# The ups and downs of biological oscillators: A comparison of time-delayed negative feedback mechanisms

**DOI:** 10.1101/2023.03.03.530971

**Authors:** Jan Rombouts, Sarah Verplaetse, Lendert Gelens

## Abstract

Many biochemical oscillators are driven by the periodic rise and fall of protein concentrations or activities. A negative feedback loop underlies such oscillations. The feedback can act on different parts of the biochemical network. Here, we mathematically compare time-delay models where the feedback affects production and degradation. We show a mathematical connection between the linear stability of the two models, and derive how both mechanisms impose different constraints on the production and degradation rates that allow oscillations. We show how oscillations are affected by the inclusion of a distributed delay, of double regulation (acting on production and degradation), and of enzymatic degradation.

## Introduction

### Biological oscillators

Periodic phenomena influence many aspects of our lives. Perhaps the most obvious examples are the repeating of the seasons and cycles of day and night. Periodicity also plays an important role on the cellular level. The 24h-rhythm that aligns with day and night is present in single cells, and many other cyclic processes govern development and survival of cells, and by extensions of tissues and whole organisms. Besides the circadian rhythms, examples include the cell cycle or the segmentation clock among numerous others (Beta and Kruse 2017).

The periodic nature of these phenomena in the cell is typically manifested by cyclic changes in the concentration or activity of different molecules. These cycles are the result of the interactions between many different genes and proteins. Despite the often bewildering complexity of protein interaction networks, many oscillatory systems rely on core network motifs to generate their oscillations (Novák and Tyson 2008).

One of these motifs is the time-delayed, nonlinear, negative feedback loop. In a negative feedback loop, a component inhibits its own activity. This property is essential for oscillations: in order to obtain periodic behavior, the system needs to be reset after one cycle. Negative feedback in itself is not sufficient for oscillations, but can also lead to stable steady states. However, feedback that acts with a sufficient time delay can result in oscillations (Casani-Galdon and Garcia-Ojalvo 2022; Ferrell, Tsai, et al. 2011; D. S. Glass et al. 2021). There are many mechanisms that can lead to time delays in biological systems. Examples include the time a signal takes to move to another location (Macnamara and Chaplain 2016; Naqib et al. 2012), or the presence of multiple intermediate steps in a system of chemical reactions (Beguerisse-Díaz et al. 2016; Korsbo and Jönsson 2020).

The study of biological oscillators has benefitted significantly from the tools of mathematical modeling. The mathematical study of (bio)chemical oscillators arguably started with the work of Lotka (1910, 1920), who showed that relatively simple chemical reactions could lead to periodic behavior. This work and later studies, such as those on the Belousov-Zhabotinsky reaction, showed that chemical systems do not have to monotonously evolve to equilibrium (Epstein and Pojman 1998; Winfree 1984). With the advent of molecular biology and the unraveling of genetic interaction networks, mathematical modeling was put to use to better understand how biochemical interactions could lead to oscillations in living systems. Here, the work of Brian Goodwin was very influential. He put forward several mathematical models to illustrate feedback mechanisms in biochemical systems, the most famous one being the three-variable Goodwin model (Goodwin 1965; Griffith 1968).

### The Goodwin model and production-inhibition oscillators

The Goodwin model consists of the equations

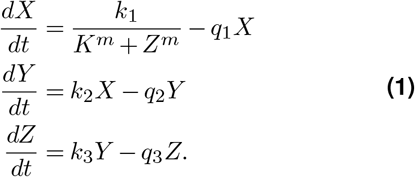

The equations model how RNA *X* is translated into protein *Y*, which activates another molecule *Z*. This finally acts as a repressor for the gene and inhibits the production of *X* (Fig. 1(a)). The inhibition is modeled by a decreasing Hill function, whose steepness is determined by the Hill exponent *m* (Fig. 1(b)). Goodwin performed simulations of the model with *m* = 1 and obtained oscillatory behavior. However, this was probably due to numerical artefacts: Griffith (1968) later showed analytically that oscillations are only possible when *m* ≥ 8. An extension of the Goodwin model includes more intermediate species (Fig. 1(a)):

**Figure 1.**
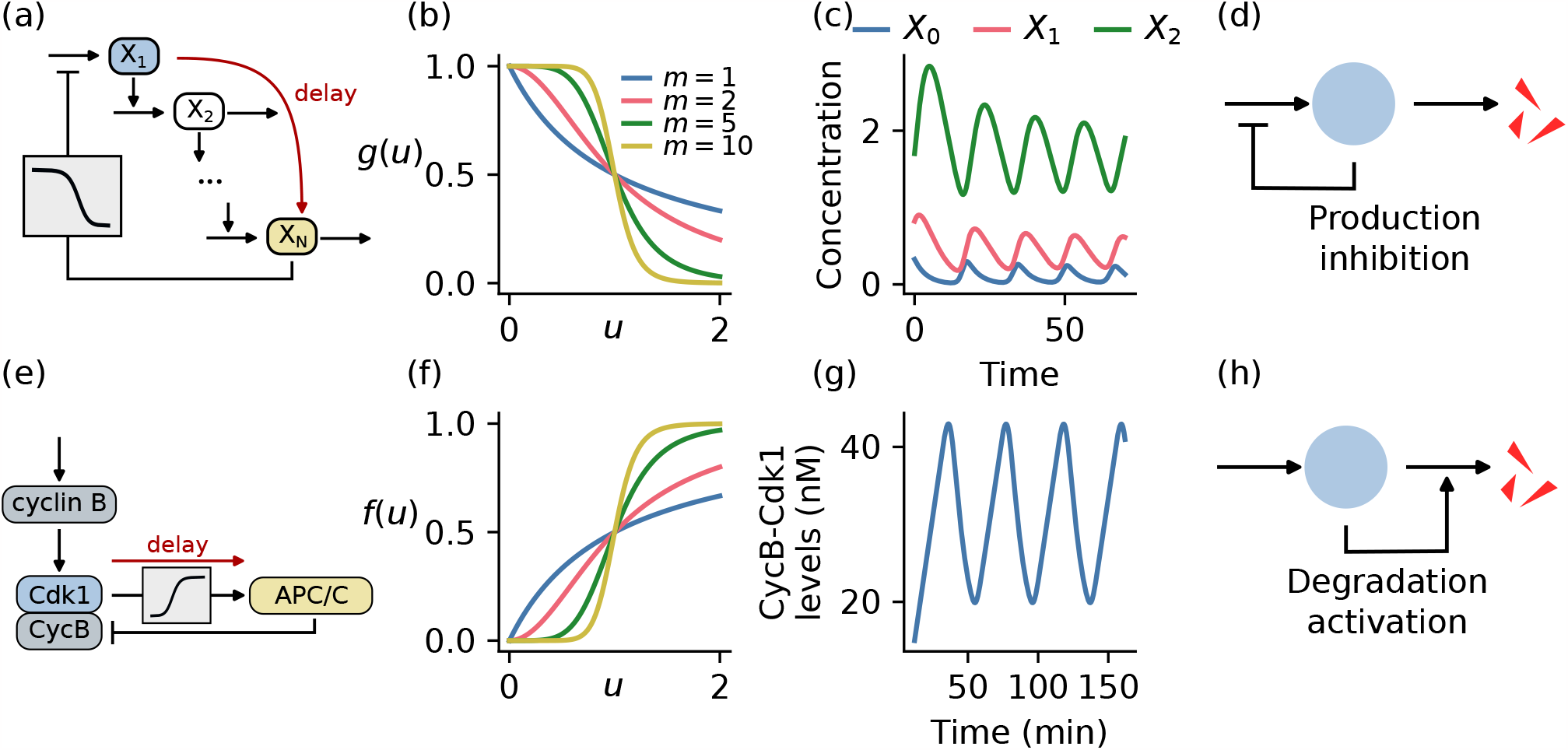
Negative feedback acting on production and degradation. (a) The multivariable Goodwin model describes a gene whose product represses its transcription. The different steps produce a time delay. (b) Inhibition can be modeled by a decreasing Hill function, with exponent *m* dictating the steepness. (c) Representative time series for the 3-variable Goodwin model. (d) The negative feedback in the Goodwin model acts on the production step. (e) Simplified model of the early embryonic cell cycle. (f) The activation of APC/C is modeled by an increasing Hill function. (f) Representative time series of Cyclin B-Cdk1 concentration in the cell cycle model. (h) In the cell cycle, the negative feedback acts through activation of degradation.

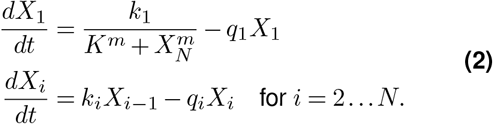

Tyson and Othmer (1978) showed that in this model, the following condition is necessary for oscillations to exist:

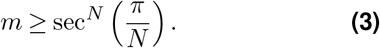

Bounds of this kind are often called *secant conditions*, for the secant function appearing in the inequality. The bound is indeed equal to 8 for *N* = 3 and asymptotically approaches 1 as *N* goes to infinity. Systems with more variables thus relax the requirement of high Hill exponents, which has been criticised as an unrealistic assumption for real biological systems (Gonze and Abou-Jaoudé 2013). The many different steps induce an effective time delay between the rise of *X*_1_ and the inhibitory effect on its own production, which promotes oscillations. An example of such oscillations in the Goodwin model are shown in Fig. 1(c), for *N* = 3 and *m* = 10. Goodwin’s model has inspired many mathematical results, from the early days (Tyson and Othmer 1978) to more recent work (e.g. Woller et al. 2014). The Goodwin model also provided an important conceptual example in whose light many other biological oscillations can be viewed. The model, or variations of it, have been used to study properties of the circadian clock (e.g. Ananthasubramaniam, Schmal, et al. 2020; Ruoff and Rensing 1996), somitogenesis (e.g. Lewis 2003; Monk 2003) and other biological cycles, and has had a profound influence on the field of biological oscillators. An excellent overview of its impact was given recently by Gonze and Ruoff (2021).

### The cell cycle model and degradation-activation oscillators

The negative feedback in Goodwin’s model acts on the production step: the final chemical species in the reaction chain inhibits the production of the first (Fig. 1(d)). Alternatively, the negative feedback could act on the degradation (Fig. 1(h)). In such a system, production is unregulated, but the degradation is induced by another molecule. A prominent biological example is the early embryonic cell cycle (Fig. 1(e)), which is driven by the periodic accumulation of cyclin proteins (Morgan 2007). Cyclin B is produced constantly, and binds to the kinase Cdk1 which is then activated. Active Cdk1 phosphorylates many different substrates, leading the cell into mitosis. One substrate of Cdk1 is a protein complex called the Anaphase-Promoting Complex / Cyclosome (APC/C). This complex targets cyclin B for degradation, leading to the inactivation of Cdk1 at mitotic exit. Here, cyclin B indirectly brings about its own destruction: a delayed negative feedback loop. Both the time delay and the nonlinear activation of APC/C have been measured (Yang and Ferrell 2013), even though the molecular mechanism of APC/C activation is not completely understood (Yamano 2019).

The many mathematical models of the early embryonic cell cycle essentially describe the feedback loop between cyclin B-Cdk1 and APC/C. In (Rombouts et al. 2018), we analyzed the equation

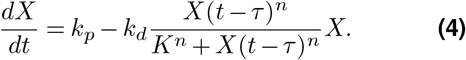

Here, *X* denotes the concentration of cyclin B-Cdk1 complexes. This concentration increases at a constant rate *k*_*p*_ and is degraded with first-order kinetics with rate *k*_*d*_. However, degradation is only active when APC/C activity is high. This activity is modeled explicitly using a Hill function (Fig. 1(f)) with argument the time-delayed term *X*(*t* − *τ*). Given suitable parameter values, this model produces cell cycle oscillations (Fig. 1(g)). Interestingly, an equivalent equation has already been used by Mackey and L. Glass (1977) to model the variation of CO_2_ levels in the blood. More generally, a feedback that activates degradation (Fig. 1(h)) can also be found in other biological systems, for example in the dynamics of the tumor repressor protein p53 in response to DNA damage (Eliaš and Macnamara 2021; Lahav et al. 2004; Lev Bar-Or et al. 2000).

### Distributed time delays

The two models — the Goodwin model and the cell cycle model — are both examples of time-delayed negative feedback loops, with a source of nonlinearity modeled by a Hill function. The time delay is explicit in the cell cycle model, but appears in an implicit way in the Goodwin model, through the introduction of multiple different steps. Indeed, in biological systems time delays often originate from multiple intermediate steps, such as phosphorylation cascades or multisite phosphorylation (Heinrich et al. 2002; Salazar and Höfer 2009). The Goodwin model with *N* steps is, in fact, equivalent to a delay equation with a distributed delay. Instead of a delay term such as *X*(*t* − *τ*), a distributed delay equation contains expressions of the form 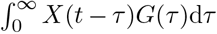 with a distribution function *G*. This means that not the value of the variable at a specific point in the past is relevant for the current time evolution, but rather its whole history, weighted by *G* (Fig. 2). As shown in the supplementary information, a linear chain of differential equations of the form

**Figure 2.**
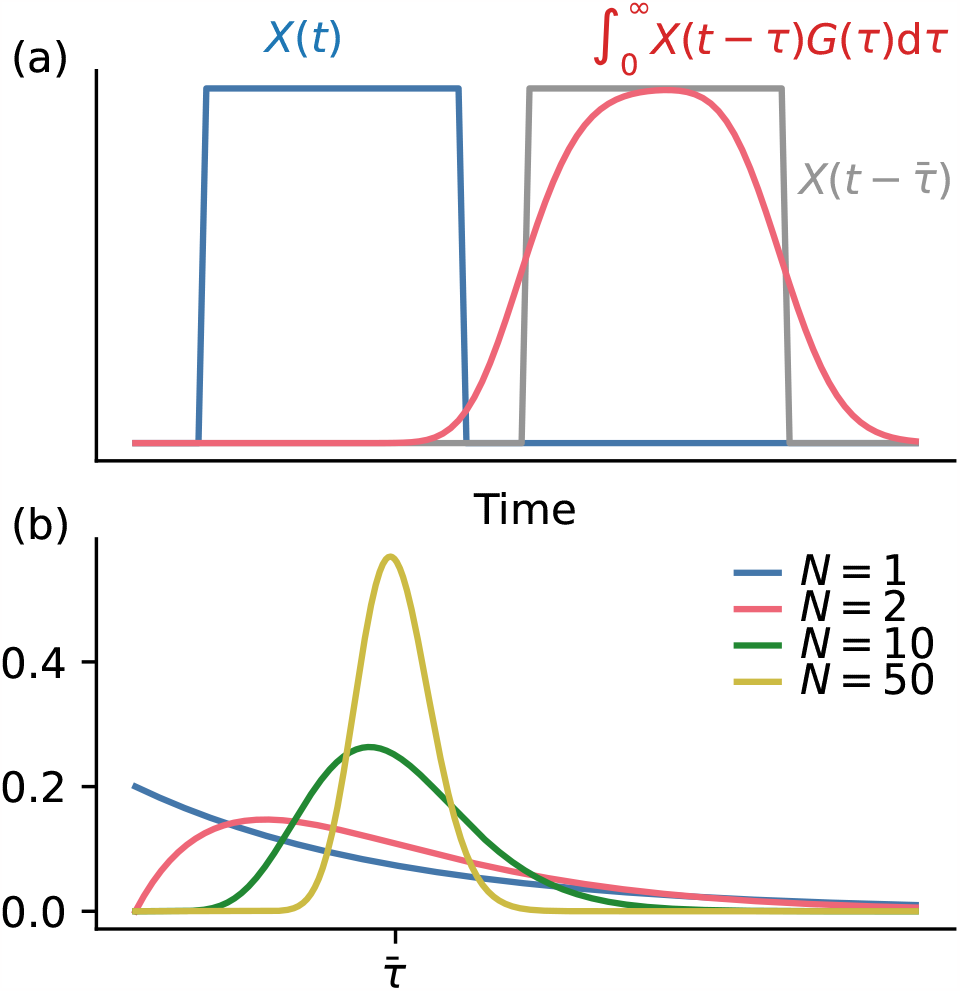
Distributed time delay. (a) A time-delayed variable with distributed delay. If the delay distribution is close to a delta distribution, the delayed variable is close to a time-shifted version of the original variable (the gray line). Here the red line was computed using a Gamma distribution with *N* = 50. (b) Gamma distributions with fixed mean but increasing values of *N* approach a delta distribution.

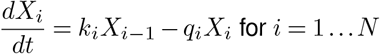

can be replaced by the expression

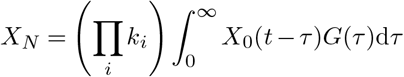

with

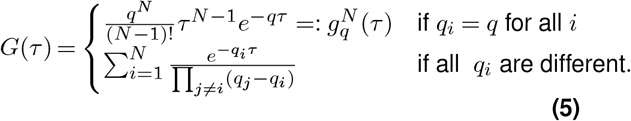

(see also (Cooke and Grossman 1982; Hinch and Schnell 2004)). In case of equal *q*_*i*_, the distribution is called the Gamma distribution. The equivalence of a Gamma-distributed delay with a system of linear ODEs is also called the *linear chain trick* (see, e.g. Smith 2010, Chapter 7). The mean of a Gamma distribution with parameters *q, N* is given by *N/q*. By keeping the mean fixed at 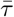 and increasing *N*, the distribution 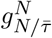 approaches a delta distribution centered at 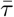 (Fig. 2b). This means that a discrete delay equation can be approximated by a long chain of ODEs, where each step happens very quickly. This equivalence also shows that the Goodwin model can be written as a single delay equation with distributed delay.

### Goal and structure of the paper

The Goodwin model on the one hand, and the cell-cycle model on the other, are examples of two different ways to implement a negative feedback loop. Each of these models has been studied in detail. Yet, a detailed mathematical study of the differences has not been done. This is what we aim to do in this paper: to provide a mathematical analysis of the most basic version of these two models, and compare their behavior. In particular, we are interested in differences between the oscillatory potential of the two models and the constraints the mechanism of feedback puts on, for example, the parameter values. This could lead to insight into why biological systems have adopted one or the other mechanism.

Our most important tools are analytical: we use linear stability analysis to determine when oscillations occur in these systems. This analysis can also be done with a graphical technique, which is sometimes more insightful than a purely algebraic one. We use this paper as an opportunity to dust off this technique, which was extensively used in the book by MacDonald (2008, originally published in 1989). We also obtain approximations for the boundaries of stability using asymptotic methods. The details of the calculations are explained in the supplementary information. For direct simulation of the delay equations, we have used the software JiTCDDE (Ansmann 2018).

Our results on the existence of oscillations have been described before, for both models: a high nonlinearity and a larger time delay lead to oscillations. However, the production and degradation rates that allow oscillations are different differ between the two. Here, we show how this difference can be explained, in fact, by a correspondence between the linear stability for the two mechanisms. If there is a destabilizing time delay for one mechanism, there is one for the other — but with an inverse value of the ratio between production and degradation rates. We interpret this result in terms of the biological rate constants and show that it also holds for Gamma-distributed time delays.

In the limit of large Hill exponents, the delay equations can be solved analytically and expressions for period and amplitude can explicitly obtained. This analysis shows that the time delay affects the amplitude significantly in one of the two mechanisms, namely the one where the feedback acts through the activation of degradation.

Finally, we shortly discuss extensions of these models. In a system where the feedback acts on both the production and degradation, we show that the oscillatory potential of both mechanisms is combined. Enzymatic degradation has a different effect on the two mechanisms.

## Results

### Model formulation

The Goodwin model and cell cycle model represent two implementations of the negative feedback loop. Both rely on a time delay and nonlinearity, which are the essential ingredients to generate oscillations. In order to study their differences, we condense these mechanisms into two delay equations that can be compared. We study the following two models, based on the Goodwin model and the cell-cycle model respectively (see Fig. 1):

### The production-inhibition (PI) model

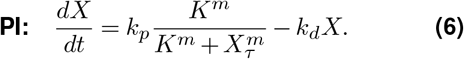

### The degradation-activation (DA) model

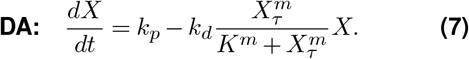

Here *X* will represent a concentration. Both models are modifications of the equation

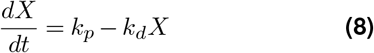

that describes constant production and first-order degradation. The PI model includes a delayed inhibition of the production rate, modeled by a decreasing Hill function (Fig. 1(a)-(d)). In the DA model, the degradation is activated with a time delay (Fig. 1(e)-(h)). With slight abuse of notation we use *X*_*τ*_ to denote either a variable with discrete delay *τ* or a distributed one with average *τ*.

In order to streamline the calculations, we nondimensionalize these equations. Setting *u* = *X/K* and *s* = *k*_*d*_*t* leads to the following equations for the production inhibition (Eq. (6)) and degradation-activation (Eq. (7)) system, respectively:

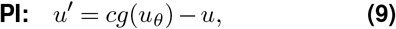

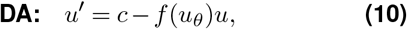

where we used the notation

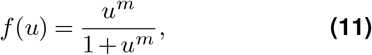

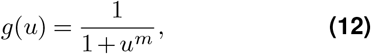

for the increasing and decreasing Hill functions and use a prime for the time derivative. Note that *f* (*u*) = 1 − *g*(*u*) = *g*(1*/u*). The only parameters in these rescaled equations are:

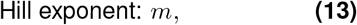

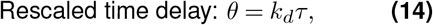

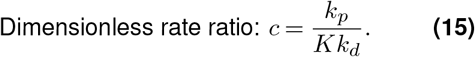

The important parameter *c* is proportional to the ratio of production and degradation rates.

### Hopf bifurcation for the system with discrete delay

In this section, we perform the Hopf bifurcation analysis of the PI and DA models (9)-(10) and compare the results. A similar analysis for the DA model can also be found in our earlier paper (Rombouts et al. 2018). For the PI model, Tyson (2002) shows the calculation. The details of the analysis can be found in the supplementary information.

First, we look at the steady states of the models. These are the values of *u* that satisfy *g*(*u*) = *u/c* for the PI model, and *f* (*u*) = *c/u* for the DA model. Plotting the two sides of these equalities (Fig. 3(a)-(b)) shows that there is exactly one positive steady state for each value of *m* and *c*. The value of the steady state increases monotonically with *c* and is independent of the time delay. To assess the stability of the steady state *ū*, we perform a linear stability analysis. For a discrete delay *u*_*θ*_ = *u*(*t* − *θ*), the linear stability analysis leads to the following equation for the eigenvalues of the systems:

**Figure 3.**
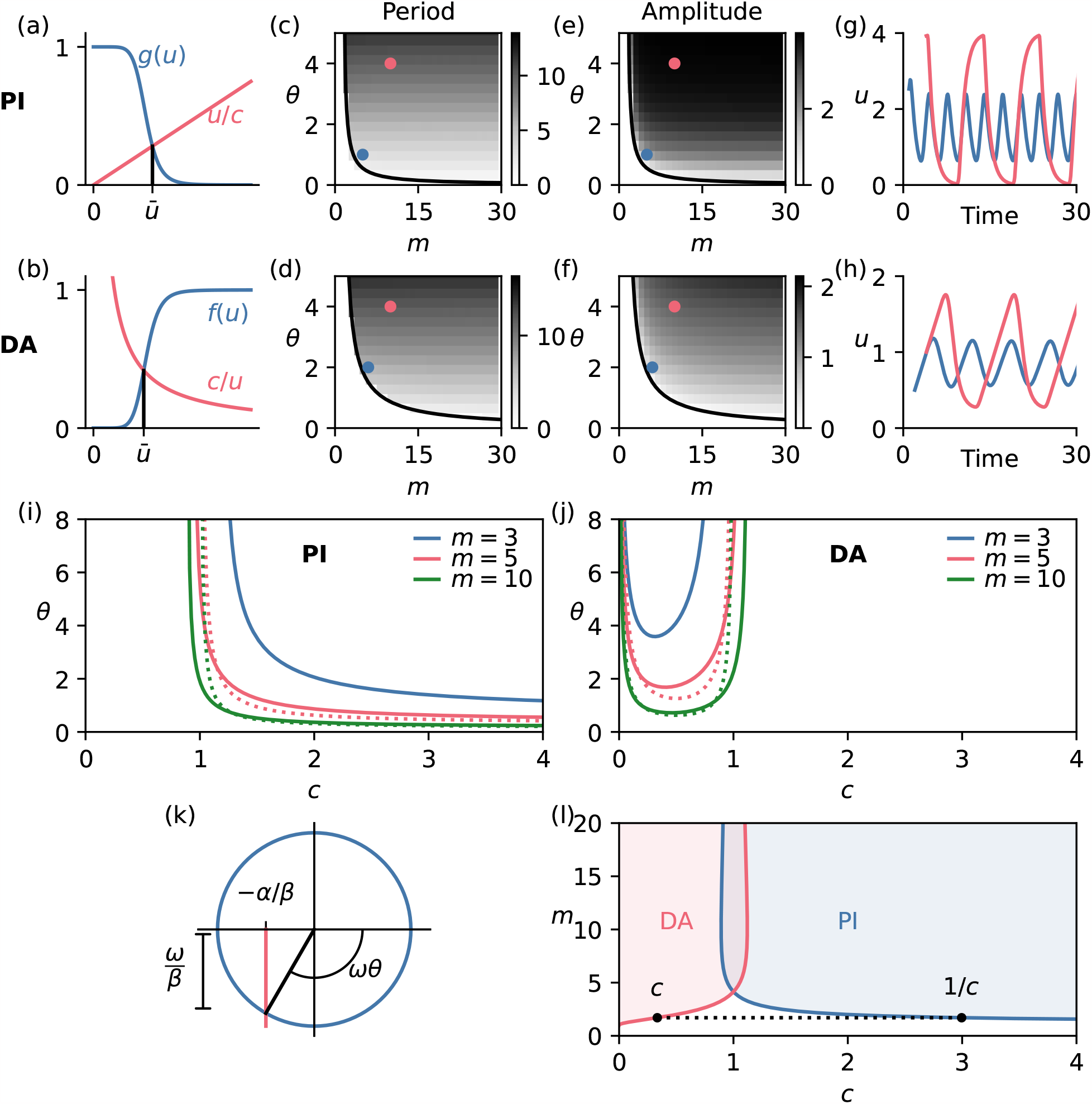
Comparison of the Hopf bifurcation for the models with discrete delay. (a) There is a unique positive steady state of the PI model, which satisfies *g*(*u*) = *u/c*. (b) The unique positive steady state of the DA model satisfies *f* (*u*) = *c/u*. (c) Oscillatory region in the (*m, θ*)-plane for the PI model, with period in color. (d) Oscillatory region in the (*m, θ*)-plane for the DA model, with period in color. (e) Oscillatory region in the (*m, θ*)-plane for the PI model, with amplitude in color. (c) Oscillatory region in the (*m, θ*)-plane for the DA model, with amplitude in color. In (c,e), we used *c* = 4 and in (d,f), *c* = 1*/*4. (g) Time series of the oscillations in the PI model, corresponding to the colored dots in panels (c,e). (h) Time series of the oscillations in the DA model, corresponding to the colored dots in panels (d,f). (i) Oscillatory region for the PI model in the (*c, θ*)-plane for different values of *m*. The dotted lines are the large *m* asymptotic expressions given in Eq. (20). (j) Oscillatory region for the DA model in the (*c, θ*)-plane for different values of *m*. The dotted lines are the large *m* asymptotic expressions given in Eq. (21). (k) Geometric interpretation of the existence of a destabilizing delay. There exists a destabilizing time delay if the circle and the vertical line intersect. (l) Regions in the (*c, m*)-plane for both models, where a destabilizing time delay exists (disregarding its numerical value). The boundaries are given by *c* = *m*(*m* − 1)^−1−1*/m*^ for the PI model and *c* = (*m* − 1)^1+1*/m*^*/m* for the DA model. The boundaries map onto each other under *c* ↔ 1*/c*.

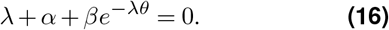

This equation is the same for both models, but *α* and *β* are different and depend on the steady state *ū*:

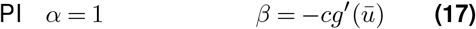

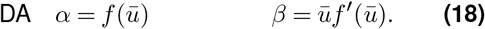

These values are always positive. For nonzero delays, the characteristic equation Eq. (16) is a transcendental equation with an infinite number of solutions. The steady state is unstable if at least one eigenvalue has positive real part. If the delay is zero, there is a single solution *λ* = − *α* − *β*. The steady state is thus stable in the absence of a delay. We next analyze whether increasing the delay can lead to oscillations through a Hopf bifurcation (see Erneux 2009; MacDonald 2008; Smith 2010). We thus look for values of the parameter values such that *λ* = *iω* is a solution of Eq. (16). Substituting this condition into Eq. (16) leads to

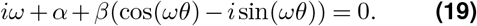

Separating real and imaginary parts and performing some algebra leads to conditions on *m, c* and *θ*, which can be used to plot regions in parameter space where the steady state is unstable (Fig. 3(c)-(f)). In these regions the model shows oscillations. For both the PI and the DA models, a sufficient delay and nonlinearity — meaning large *m* — are needed to obtain oscillations. There is a trade-off: if the Hill exponent *m* is larger, a smaller delay is sufficient for oscillations to occur. An asymptotic analysis, worked out in the supplementary information, reveals that the critical time delay scales as 1*/m* for large *m* (see also Supp. Fig. S1,S2). In particular,

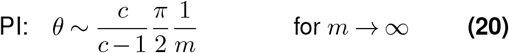

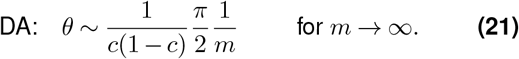

The oscillatory regions in the (*c, θ*)-plane are different between the two mechanisms: for the PI model, a large value of *c* promotes oscillations, and oscillations are impossible when *c* is roughly less than 1 (Fig. 3(i)). For the DA model, oscillations are only possible when *c* is between zero and (approximately) 1 (Fig. 3(j)).

More insight into the existence of a destabilizing time delay can be gained by considering the Hopf bifurcation condition in a geometric way (MacDonald 2008). We rewrite Eq. (19) as

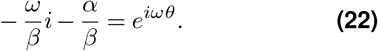

A Hopf bifurcation is possible if there exist values of *ω* and *θ* such that this equality holds. When freely varying *ω* and *θ*, the left hand side describes a vertical line in the bottom half of the complex plane. This line intersects the real axis at −*α/β*, which is less than zero since *α* and *β* are positive. The right hand side describes the unit circle. If the line and the circle intersect, there is a time delay that destabilizes the steady state (Fig. 3(k)). Whether there is an intersection only depends on the value of *α/β*, which in turn only depends on the parameters *m* and *c*. In particular, there is an intersection if *α/β <* 1. Fig. 3(l) shows the region in (*c, m*)-plane where this condition is satisfied. The boundaries have an analytical expression (see the supplementary information). For values of *c* and *m* above the lines, there exists a destabilizing time delay. Remarkably, there is a correspondence between these regions for the PI and the DA model: they map onto each other under the correspondence *c* ↔ 1*/c*. Concretely: if for the PI model with parameters (*c, m*) there exists a critical time delay which makes the steady state unstable, then there also exists such a time delay for the DA model with parameters (1*/c, m*). This correspondence is essentially due to the fact that the geometric picture for the corresponding models is the same. We show the correspondence now algebraically, by rewriting the condition *α/β <* 1.

We use Eqs. (17) and (18), together with the condition for the steady state, to rewrite this condition for both models:

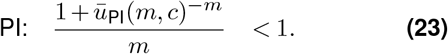

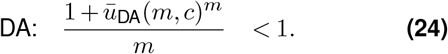

We have explicitly written the steady states of each models as a function of *m* and *c* here. These conditions cannot be satisfied for *m* = 1 since the steady state *ū* is larger than zero. This shows that for both models, oscillations are impossible for *m* = 1. The conditions have the same form, except that for the PI model the numerator contains *ū*^−*m*^ and for the DA model it is *ū*^*m*^. To show that these conditions are the same for both models, when using *c* for one and 1*/c* for the other, we now show that

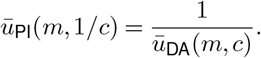

We have that *ū*_PI_(*m*, 1*/c*) is the single positive root of the equation

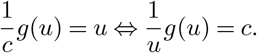

As *g*(*u*) = *f* (1*/u*), the equation becomes

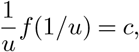

which states that 1*/u* is the unique positive steady state of the DA model with parameter *c*.

These calculations show that a Hopf bifurcation occurs at a critical time delay in the PI model if and only if it happens in the DA model, with inverse value of *c*. However, the actual values of the critical time delays need not be the same. From the geometrical picture and Eq. (22), we know that the coordinates in the complex plane of the intersection point are the same for both models (taking into account the *c* ← 1*/c* correspondence), such that

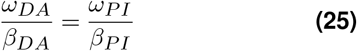

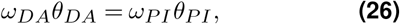

from which we derive

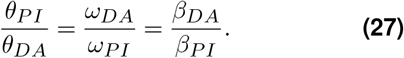

By working this out, and using the relation between the steady states of both models, we find

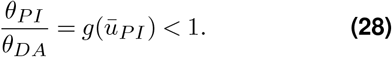

This entails that the delay needed to destabilize the PI model is smaller than the delay for the corresponding (that is, inverse *c*-value) DA model. The difference can be very large: the regime diagram for the PI model in the (*c, θ*) plane (Fig. 3(i)) shows that if *c* → ∞, the critical time delay approaches a finite value, which can be computed as (see supplementary information)

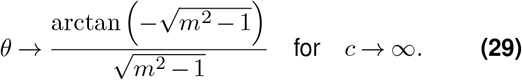

For the DA model model, however, the critical time delay goes to infinity as *c* → 0 (Fig. 3(j)).

The difference between both mechanisms thus mainly lies in the role of the parameter *c*, which we defined as *c* = *k*_*p*_*/*(*Kk*_*d*_). The way the negative feedback acts constrains the production and degradation rates that allow oscillations. For the PI model, the production rate can be very large without compromising the potential for oscillations, but a low production rate prohibits oscillations. The inverse holds for the DA model. The value of *c* = 1 corresponds to *k*_*p*_*/k*_*d*_ = *K*. The ratio *k*_*p*_*/k*_*d*_ is the steady state of the production-degradation system without any regulation: *X*′ = *k*_*d*_ − *k*_*d*_*X*. The condition on *c* thus corresponds to comparing the steady state of the unregulated system with the threshold of the activation/inhibition functions.

### Hopf bifurcation for distributed delay

A discrete delay is somewhat artificial, and unlikely to represent biological delay mechanisms accurately. A more realistic description uses a distributed delay. A convenient delay distribution is the Gamma distribution, as discussed above (see Fig. 2). We now consider the dynamics of a system with a Gamma-distributed delay. We fix the mean delay 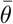 and change the variance by varying the parameter *N*. Increasing *N* corresponds to a sharper distribution (Fig. 2b). As detailed in the supplementary information, the characteristic equation analogous to Eq. (16) in the case of a Gamma-distributed delay with average delay 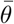 and parameter *N* is

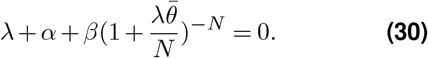

The parameters *α* and *β* are the same as previously for both models. This equation can be rewritten as a polynomial equation, echoing the equivalence between a system with Gamma-distributed delay and a set of ODEs. Performing the Hopf ansatz *λ* = *iω* and rewriting yields

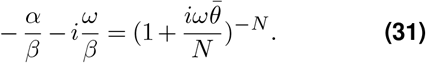

We apply the same geometric reasoning to this equation as to Eq. (22). The left-hand side describes a vertical line in the lower half of the complex plane, whose position is determined by the ratio *α/β*. The right hand side describes a spiral parameterized by 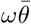 whose shape depends on *N* (Fig. 4a and Supp. Fig. S3). For *N* = 1, the curve is a half-circle with only positive real part. This shows that for *N* = 1, an instability cannot occur. For *N* = 2, the curve is the lower half of a cardioid. In general, a polar form of the spiral is given by

**Figure 4.**
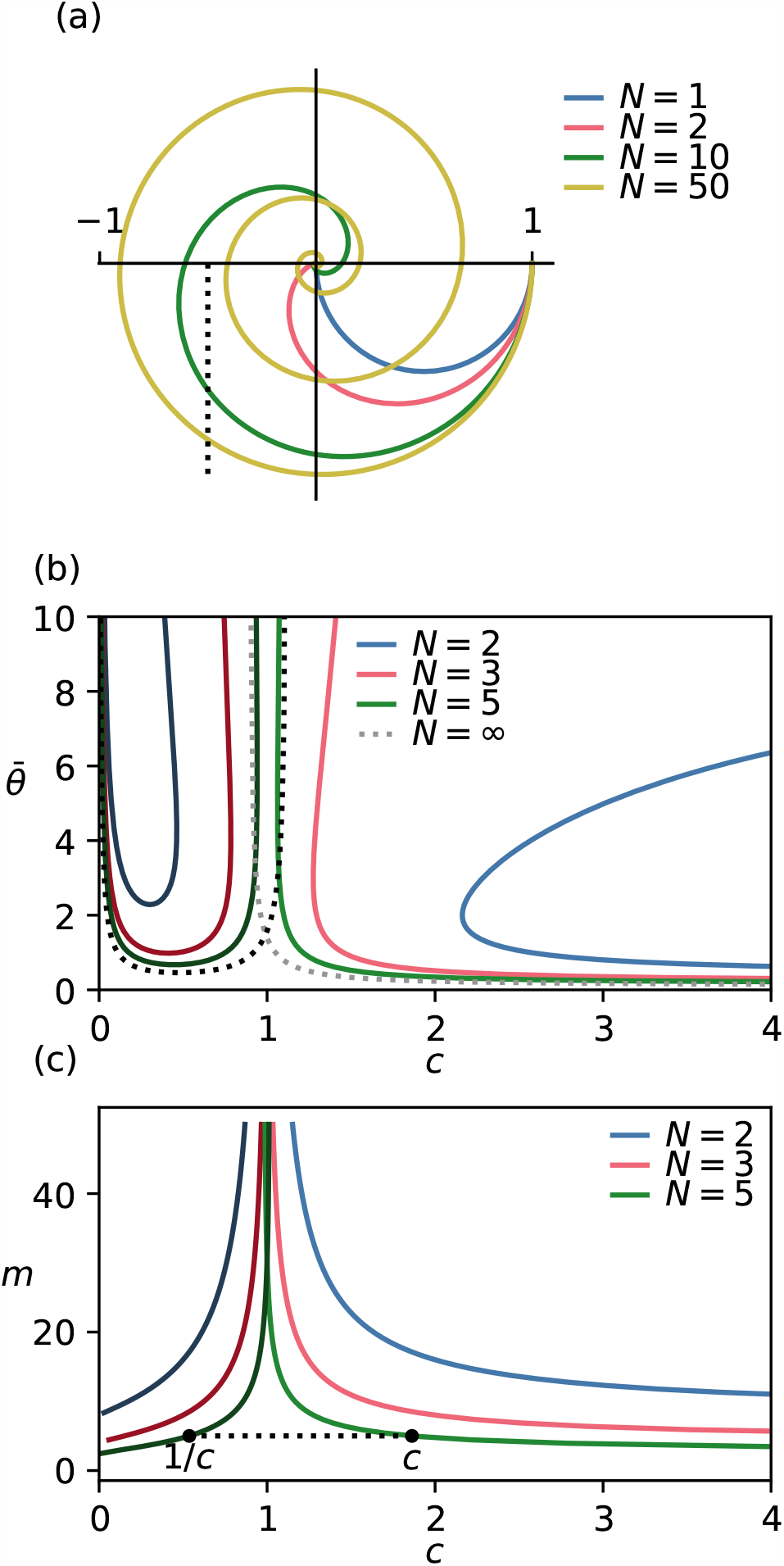
Hopf bifurcation for systems with Gamma-distributed delay. (a) Spirals corresponding to Gamma distributions with increasing *N*. A critical time delay exist if the spiral intersects with the vertical line at *− α/β*. (b) Stability boundaries in the 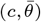 plane for different values of *N*, for both models. Darker color is the DA model, lighter color is the PI model. Dashed lines are the stability boundaries for the models with discrete time delay. We used *m* = 15. (c) Region in the (*c, m*) plane where a destabilizing average delay exist, for different values of *N*. Darker color: DA model, lighter color: PI model. For each *N*, the dark and light curve map onto each other through *c ↔* 1*/c*.

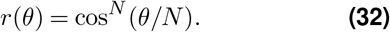

This is also called a sinusoidal spiral. As *N* become large, the part of the spiral that lies in the bottom half of the complex plane approaches a semicircle: the curve for the case of a discrete time delay.

This geometrical description directly shows that also for Gamma-distributed delays, there is a correspondence between the PI model and the DA model when it comes to the possibility of a Hopf bifurcation. Given *N*, there exists an *average* delay that destabilizes the steady state for one mechanism if and only if it exists for the other mechanism with inverse *c*. Fig. 4c shows this graphically: the lines demarcate the regions in (*c, m*)- space where a critical time delay exist. These lines map onto each other for the two models under the transformation *c* ↔ 1*/c*. There is an analytical formula for the critical *c* as function of *m* given in the supplementary information. It can be derived by solving − *α/β* = *Q*(*N*), where *Q*(*N*) is the minimal real part of the spiral.

There are values of *α/β* for which the vertical intersects the spiral twice (Fig. 4a). In such a situation, there are two critical average time delays. When the time delay increases past the first value, the steady state becomes unstable and there are oscillations. When the time delay increases even more and crosses the second critical value, the steady state regains stability. This behavior is not seen for discrete delays. The restabilization is also clear from the regime diagrams in Fig. 4b: for certain values of *c*, there are two critical time delays. A further discussion of this phenomenon can be found in MacDonald’s book (MacDonald 2008).

### Analytical solution for the limit of large Hill exponents

The Hill functions we use to model the feedback approach a step function in the limit of large Hill exponents (see also Fig. 1(b,f)). In the system with discrete delay, this can be exploited to obtain an analytically solvable system (Erneux 2009; Mackey 1997). For example, for *m* → ∞ the PI system reduces to

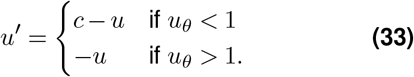

This equation is piecewise linear. Solving on each piece and gluing together the results gives an expression for *u*(*t*). The time series are shown in Fig. 5(a,b) and Supp. Fig. S5. The details of the calculation are given in the supplementary information. Of interest to us are the resulting explicit expressions for period *P* and amplitude *A*. For the model with inhibition of production (PI) we find

**Figure 5.**
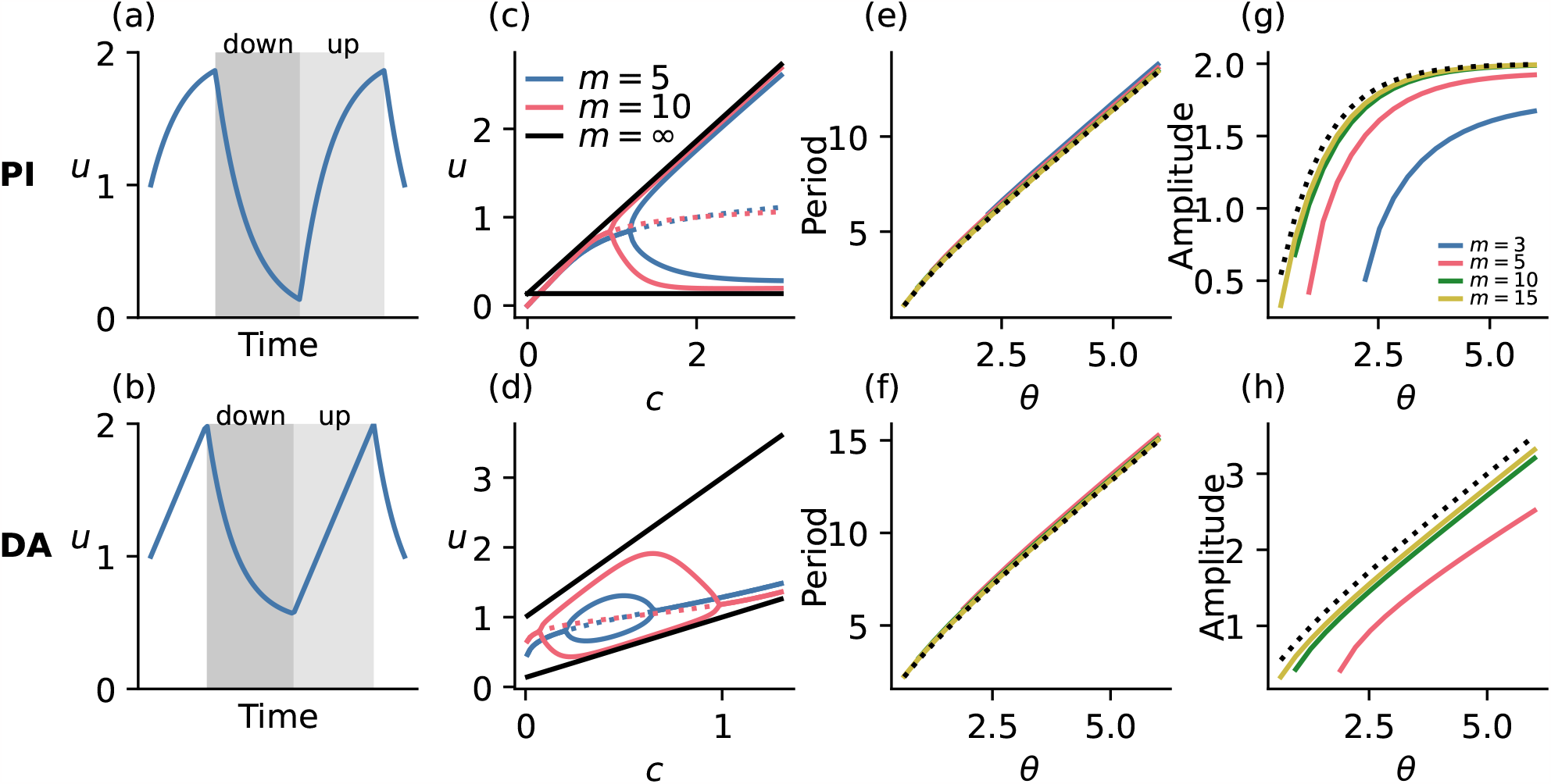
The models with *m → ∞* are analytically solvable, and provide good approximations for period and amplitude. Top row: results for the PI model, bottom row: results for the DA model. (a,b) Time series of the models for *m → ∞*. The equation can be solved exactly on the up and down parts. (c, d) One-parameter bifurcation diagram: maximal and minimal value of *u* as function of *c*. The black lines are analytical expressions obtained in the limit *m → ∞*. The dashed line is the steady state which is unstable in the case of oscillations. (e,f) Period as function of time delay for different values of *m*, with the analytical prediction as dashed line. Note that the lines for finite *m* do not start at *θ* = 0, but only at the critical *θ* defined by the Hopf bifurcation. (g,h) Amplitude as function of time delay for different values of *m*, compared to the analytical prediction for *m → ∞* (dashed). For panels (e) and (g), *c* = 2 and for panels (f) and (h), *c* = 1*/*2.

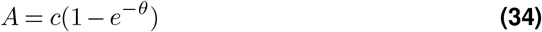

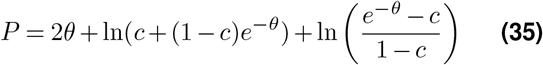

and for the model with activation of degradation (DA) we have

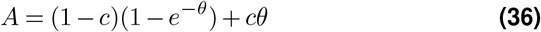

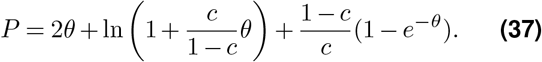

Important to note is that we are working with scaled units. The unscaled amplitude is multiplied by *K* and period by 1*/k*_*d*_. Moreover, the formulas are only valid for *c >* 1 in the case of the PI model and for 0 *< c <* 1 in the case of the DA model. These formulae give a good approximation of the amplitude and a very good approximation of the period, even for not-so-large values of *m* (Fig. 5(c-h). We can use these formulae to analyze differences between the oscillators far from the oscillatory instability.

For large time delays, the amplitude depends on the time delay for the DA model, but is independent of it for the PI model. The fact that amplitude does not depend strongly on the time delay seems to be more generally true for oscillators based on delayed inhibition of production (Jörg 2017). Since *c <* 1 for the DA model, we see that the amplitude *A* satisfies *A <* 1 − *e*^*−θ*^ + *θ* for this model. So, very large amplitude oscillations in the DA model are only possible for large time delays. For both models, the period contains a term 2*θ*: the time delay contributes significantly to the period. One interesting question, inspired by the paper by Mather et al. (2009), is whether oscillations are possible that have a much longer period than the time delay. The answer is yes in principle, for both models: for the PI model, the period diverges for *c* → ∞ and *c* → 1. For the DA model, *P* goes to infinity for *c* → 0 and *c* → 1. Large periods are thus possible if the production and degradation rates are either very imbalanced, or when *k*_*p*_*/k*_*d*_ ≈ *K*. However, the divergence of *P* is logarithmic, so very slow, except in the case of the DA model with *c* → 0. This suggests that between these mechanisms, the most efficient waym of obtaining periods much larger than the time delay is in the DA model, with production rate much smaller than degradation rate.

### Enzymatic degradation

We finally consider two variations of the model. First, we study a variation of the model where the rate of degradation is limited by the availability of an enzyme. The possible importance of saturating degradation was already present in Goodwin’s early work (Gonze and Ruoff 2021; Goodwin 1965). Moreover, it has been suggested that including a enzymatic degradation in one of the variables can alleviate the necessity of high Hill exponents in the classic Goodwin model (Tyson 2002).

For the PI model, changing first-order into Michaelis-Menten-like kinetics for the degradation yields

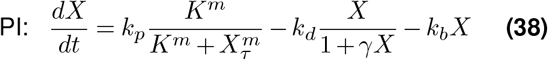

For the DA model we have

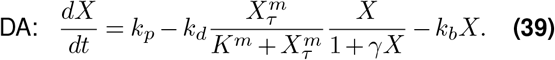

The degradation is no longer given by *k*_*d*_*X* but by *k*_*d*_*X/*(1 + *γX*) + *k*_*b*_*X*. We have included a basal first-order degradation at rate *k*_*b*_ in order to avoid the unbiological situation in which there exist no steady states, which may happen for *k*_*b*_ = 0 in the DA model if the maximal enzymatic degradation rate is smaller than the production rate. In order to stay close to the models of the previous sections we consider *k*_*b*_ ≪ *k*_*d*_. We also restrict ourselves to the discrete delay case.

For small values of *X*, the term 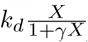 is approxi-mately equal to *k*_*d*_*X*. This corresponds to the situation where there is enough enzyme such that the degradation is proportional to the concentration *X*. For larger values of *X*, the new degradation term saturates and approaches *k*_*d*_*/γ*. Performing the same rescaling as before leads to the two equations

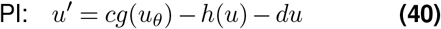

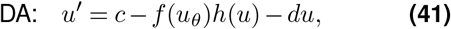

with *f* and *g* defined above, *c* = *k*_*p*_*/*(*k*_*d*_*K*), *d* = *k*_*b*_*/k*_*d*_ and

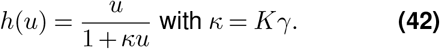

For *d* = *κ* = 0, these equations reduce the ones from the previous sections. It is thus instructive to start by looking at the behavior of these models for relatively small *κ* and *d*, to see how the introduction of enzymatic degradation changes the dynamics relative to the basic models. The linear stability analysis of this system is given in the supplementary information. We find differences in the PI and DA mechanisms.

For the PI model, a non-zero but relatively small value of *κ* increases the range of both *m* and *c* for which oscillations exist (Fig. 6(a,c,e)), but the effect is partially reversed when *κ* becomes large (Fig. 6(g)). Remarkably, oscillations are also possible for values of *m* smaller than one. However, the time delay needed for the steady state to become unstable typically increases. For the DA model, on the other hand, a *κ* slightly larger than zero leads to a smaller region in the parameter planes where oscillations exist (Fig. 6(b,d)). It seems, however, that it is mainly the time delay which needs to be much larger, since the effect in the (*c, m*)-plane is more mixed: increasing *κ* leads to oscillations for lower *m*, but also for a smaller range of *c*, which is also the trend for much larger values of *κ*. As before, we can plot the oscillatory regions in the (*c, m*)-planes (Fig. 6(e-h)). The regions in this plane indicate values of *c* and *m* for which a destabilizing delay exist — but it says nothing about how large this delay should be.

**Figure 6.**
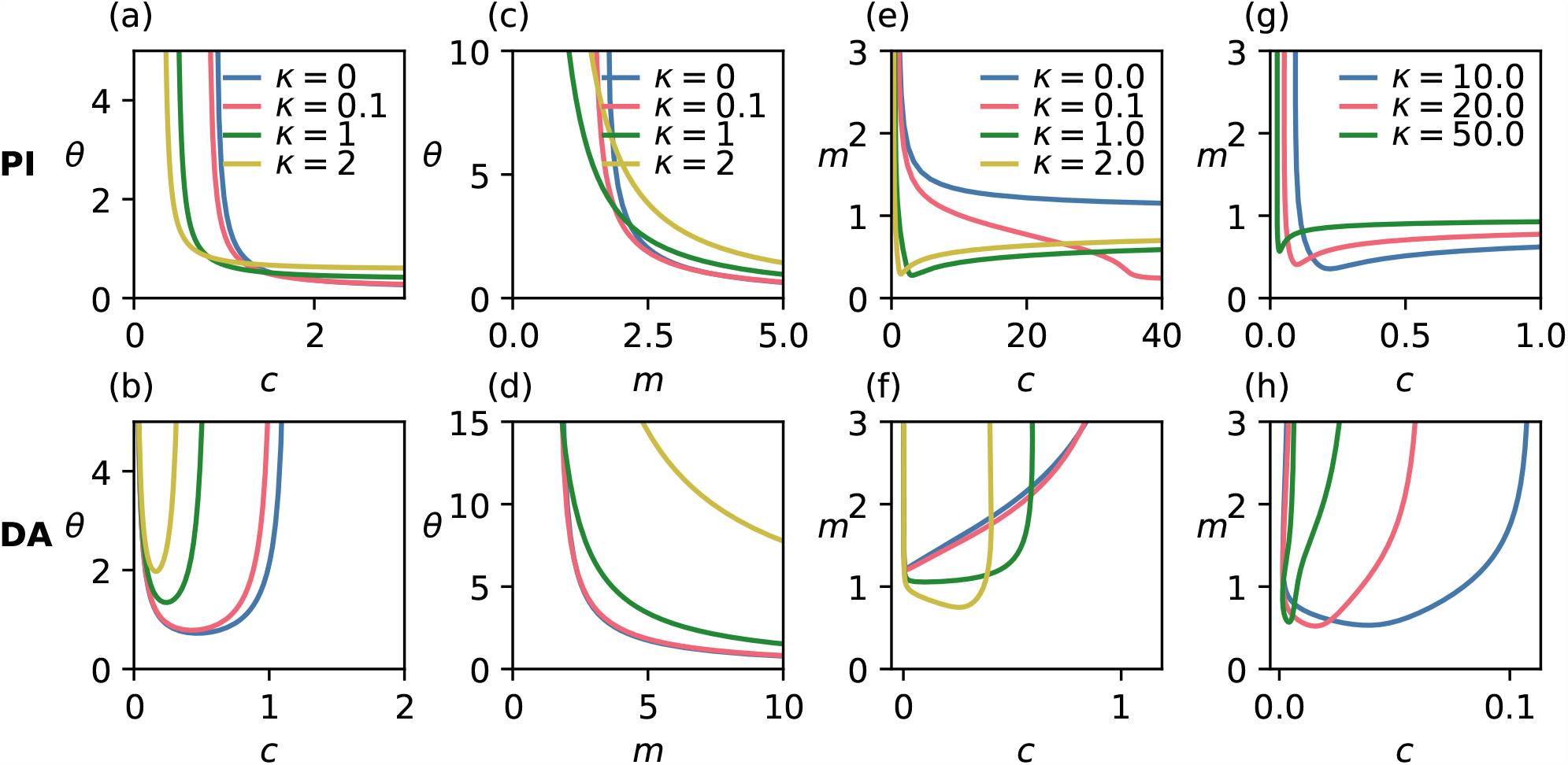
Enzymatic degradation has a different effect on both mechanisms and generally can lead to oscillations for smaller. *m*. All panels show stability boundaries defined by the Hopf bifurcation criterion. Top row: PI model, bottom row: DA model. For (a-d), the steady state is unstable and oscillations exist for parameter values above the lines. For (e-h), above the line there exists a time delay which destabilizes the steady state, and below delay it is stable for all delay values. See also Supp. Figs. S6,S7. For all panels, *d* = 0.01. Panel (a,b): *m* = 10, panel (c): *c* = 3, panel (d): *c* = 1*/*3.

The results for small *κ* tell us something about the biological system where the enzyme is only limiting for large concentrations of *X*, i.e. where there is a large availability of enzyme. A more biologically relevant situation is when the enzyme is limited and the degradation saturates quickly, corresponding to large values of *κ*. As shown in Fig. 6(g,h), oscillations in this regime can appear even for small values of *m*, but they require small values of *c* too. Moreover, the stability lines can develop different bumps and wiggles, indicating nontrivial stability changes (see also Supp. Figs. S6,S7). Finally, in this system it might be possible to have both oscillations and a stable steady state for the same set of parameters, such that the stability boundaries do not correspond with the existence of oscillations per se (see supplementary information). We leave the more detailed analysis of the models with enzymatic degradation to future work.

### Double regulation

What if the PI and DA mechanisms are combined into a system where both production and degradation are regulated? Such a system would be modeled by the equation

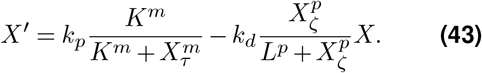

In principle, the thresholds, Hill exponents and delays for both terms can be different. Here, we do not provide a detailed analysis for all possible parameters. Instead, we aim to illustrate some properties of this model by analyzing the case of equal parameters: *m* = *p, K* = *L* and *τ* = *ζ*. With this assumption, rescaling leads to

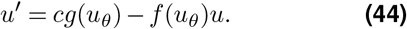

The stability analysis is done as before, and reveals that this model combines the oscillatory potential of PI and DA models: oscillations are possible for any positive value of *c* (Fig. 7a). This is also true for the model with a Gamma-distributed delay: whether a destabilizing time delay exists for a given *N* only depends on *m* and not on *c*, because the value of *α/β* is exactly equal to 1*/m* for this model. For this model, it is straightforward to compute the values of *N* for which there is a destabilizing delay, and for which values of *N* and *m* the steady state is restabilized for increasing time delays (Fig. 7b). In the supplementary information we explain that a restabilization occurs for values of *m, N* such that

**Figure 7.**
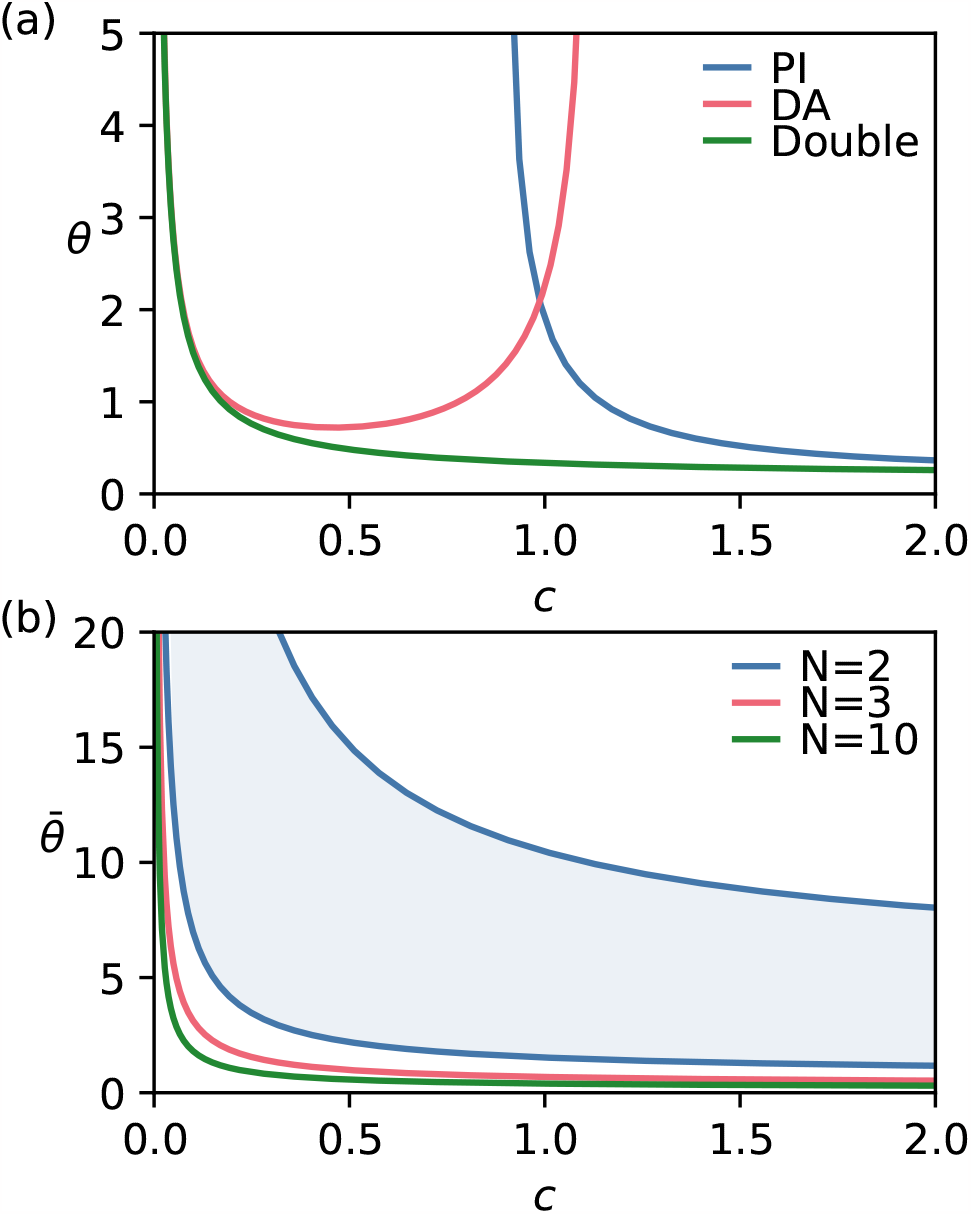
Oscillations in the system with double regulation. (a) Regime diagram for different mechanisms, defined by the Hopf bifurcation line. Oscillations exist for parameter values above the curves. (b) Boundary of the oscillatory region with Gamma-distributed delay for different *N*. For *N* = 2, only the parameter values between the two blue curves lead to an unstable steady state. Both diagrams were computed with *m* = 10.

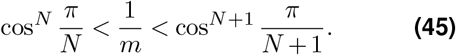

For *N* = 2, this means that there is a restabilization for all *m >* 8 and for *N* = 3, for *m* between 4 and 8. For *N <* 2, no oscillations are possible. The range of values of *m* for which restabilization is possible becomes smaller and smaller as *N* increases (see also Supp. Fig. S4).

## Discussion

Biological oscillations are typically characterized by cyclic changes in the concentration or activity of different molecules. Whereas activities can be modulated, for example, by posttranslational modifications, concentration changes are often generated by regulated production and degradation. In this work, we have compared two mechanisms that are key to generate such production-degradation oscillations. Whereas the high-level logic of both mechanisms is the same — a time-delayed negative feedback loop — the molecular implementation is different. In one oscillator, the feedback acts to inhibit production (the PI model), while in the other mechanism it acts to activate degradation (the DA model). Inspired by the Goodwin model and its many applications on one hand and the cell division cycle on the other, we studied these oscillators in their most simple version: a single delay differential equation with nonlinearity modeled by a Hill function.

Our mathematical analysis revealed a connection between the two oscillators: if oscillations exists (for large enough time delay) for one model, then this is also the case for the other model, but with the inverse of the parameter *c*. The result is valid both for discrete and Gamma-distributed delays. It implies that both mechanisms impose different constraints on the ratio of production to degradation rates: if the feedback is implemented through inhibition of production, the system has the potential to oscillate when *c* is large, i.e. production rate is much higher than degradation rate, and the inverse holds for the DA model. This connection does not address the magnitude of the time delays required to obtain oscillations. This differs for both models: for large *c*, the required time delay for the PI model approaches a constant value, whereas it diverges for small *c* for the DA model.

We also derived asymptotic approximations which — to our knowledge — have not been described in the literature, such as 1*/m* scaling of critical time delay as function of Hill exponent. Analytical results can also be obtained far from the threshold of instability, by exploiting the fact that the Hill function approaches a step function — a trick that has been used many times before. The calculation yields explicit formulae for amplitude and period, revealing differences between the two mechanisms. The amplitude continues to increase with larger time delays in the DA model, while it saturates for the PI model. This is consistent with previous findings that the oscillation amplitude in PI models does not depend strongly on the delay (Jörg 2017). The main contribution to the oscillation period is given by twice the delay. Between the two mechanisms considered, the easiest way to obtain much larger periods than twice the delay time is by having the production rate much smaller than degradation rate in the DA model.

Finally, we showed how two extensions — double regulation and enzymatic degradation — modify the oscillatory potential of both oscillators. Since these extra mechanisms introduce many different parameters, we only investigated a subset of the possible behavior. We leave a more in-depth analysis of the dynamics of these systems for later work.

Some of our results, such as the exact correspondence of oscillatory potential with the *c* ↔ 1*/c* correspondence, rely on specific properties of, for example, the Hill functions. Nevertheless, we believe that the general messages are likely to be more generally applicable. The two basic mechanisms we studied here underlie many real biological oscillators, and our results for the simplified models can be used to guide the study and interpretation of more complex models. Confirming the generic results in a more molecularly detailed model would be an interesting follow-up. In particular, the time delay and the Hill function are high-level descriptions of what is happening at the molecular level. There are many different ways in which an ultrasensitive response can be generated (see Ferrell and Ha (2014a,b) for an overview and Jeong et al. (2022) for an example of recent work on this) and the same holds for the time delay. We have assumed these to be independent, but delay and ultrasensitivity could also be due to one single molecular mechanism, connecting these two parameters.

Real biological oscillators can be much more complicated than a single negative feedback loop (see e.g. Novák and Tyson 2008; Purcell et al. 2010). In many real systems, the negative feedback is complemented by positive feedback loops. These can make oscillations more robust and tunable (Ananthasubramaniam and Herzel 2014; Tsai et al. 2008). In a sense, one type of positive feedback is included in our model through the ultrasensitive Hill-type response curve, as such nonlinear switch-like responses can be the result of positive feedback. However, positive feedback can also generate bistability, which we have not considered here. The combination of negative feedback with a single (Novák and Tyson 2008; Tsai et al. 2008) or multiple (De Boeck et al. 2021; Parra-Rivas et al. 2023) bistable switches can lead to robust relaxation oscillations. It would be interesting to systematically investigate the difference between the PI and DA mechanisms in models that also incorporate such bistable switches.

The importance of how a negative feedback is implemented has recently also been put forward by Agrahar and Rust (2022, preprint). In that study, the authors consider different implementations based on regulation of production and degradation, but they did not consider explicit time delays. They use a computational approach: by sampling parameter values, they determine which oscillator implementations are most likely to yield oscillations. Some of our current results echo theirs. For example: Agrahar and Rust (2022) also found that dual regulation increases oscillatory potential. Other recent studies have considered the question of robustness, typically by large-scale sampling of parameter values. Li et al. (2017) find that adding small auxiliary motifs to core topologies may enhance the robustness of the oscillations. Furthermore, they discuss how an evolutionary process could have generated such topologies. Besides insights into the evolutionary origin of regulatory mechanisms, such studies can also be important to guide the design of synthetic oscillators (e.g. Woods et al. 2016).

Our current results, which focus on in-depth mathematical analysis of a basic motif, complement the cited studies that use a more computational approach. These studies also suggest useful considerations to develop further mathematically. Whereas now, we only considered differences in the parameter ranges that can lead to oscillations, the PI and DA models may also show differences in other aspects. For example, a study of the energetic cost of the mechanisms would be a useful next step. An additional factor which may differ between mechanisms is whether it can be entrained by external cues (Jiménez et al. 2022). For example, for the circadian clock it is important that it can be entrained to the 24h rhythm of day and night. Whether this requirement imposes conditions on the negative feedback loop, and in particular whether it favours a DA or PI mechanism, is also an interesting question.

Finally, an important aspect we did not consider in our study is the role of stochasticity. This has recently been investigated by Negrete et al. (2021) for a model akin to our PI oscillator. The authors make use of the approximation for *m* → ∞ to derive statistical distributions for the extrema and rise and fall time of the oscillator. They show that experimental data from the mammalian circadian clock and the zebrafish segmentation clock can be well described by this model. There could be differences in those statistical distributions between PI and DA model — which are to be determined.

The study of biological oscillators is arguably one of the oldest topics in mathematical biology. Yet, the many recent studies illustrate that the subject is very much alive (see Tyson, Csikasz-Nagy, et al. 2022, for an overview). Great progress is made using computational methods, and more and more by incorporating real biological data. Nevertheless, our current study shows that there are still things to be learned by analyzing the most simple mathematical models.

