## Supplemental Information and Figures for "The ups and downs of biological oscillators: A comparison of time-delayed negative feedback mechanisms"

### Supplementary Information

Jan Rombouts, Sarah Verplaetse, Lendert Gelens

March 3, 2023

#### 1 The Goodwin system with different rates is equivalent to a distributed delay equation

Here we show that the system

$$\begin{aligned} \frac{dX_1}{dt} &= \frac{k_1}{K^n + X_N^n} - q_1 X_1 \\ \frac{dX_i}{dt} &= k_i X_{i-1} - q_i X_i \quad \text{for } i = 2 \dots N. \end{aligned} \tag{1}$$

is equivalent to an equation of the form

$$\frac{dX}{dt} = \frac{k_1}{K^n + X_\tau^n} - q_1 X, \tag{2}$$

where

$$X_\tau = \int_0^\infty X(t-\tau) g(\tau) \mathrm{d}\tau, \tag{3}$$

and $g$ has the form

$$g(t) = \sum_{i=2}^{N} \frac{k_i}{\prod_{j \neq i} (q_j - q_i)} e^{-q_i t} \tag{4}$$

We assume for simplicity that all the $q_i$ are different, but a general expression can be derived that also holds for equal $q_i$. If all the $q_i$ are equal, the same derivation yields a Gamma distribution. A proof by induction of the equivalence can be found in the paper by Hinch and Schnell (2004). Here, we show it directly using the Laplace transform. We take the Laplace transform of the second line of Eq. (1) to find

$$s\hat{X}_i = k_i \hat{X}_{i-1} - q_i \hat{X}_i,$$

where

$$\hat{X}_i = \mathcal{L}(X_i) = \int_0^\infty e^{-st} X_i(t) \mathrm{d}t$$

is the Laplace transform of $X_i$. We rewrite as

$$\hat{X}_i = \frac{k_i}{s+q_i} \hat{X}_{i-1},$$

which can be iterated to yield

$$\hat{X}_N = \prod_{i=2}^{N} \frac{k_i}{s+q_i} \hat{X}_1. \tag{5}$$

The latter equation has the form

$$\hat{X}_N = \hat{g} \hat{X}_1, \tag{6}$$

where $\hat{g}$ will be the Laplace transform of the distribution function we are looking for. Indeed, from Eq. (6), by taking the inverse Laplace transform we find

$$X_N(t) = \int_0^\infty X_1(t-t') g(t') \mathrm{d}t', \tag{7}$$

where we used the property that convolutions in the time domain correspond to multiplication in the Laplace domain. We now need to show that the inverse transform of the function $\hat{g}(s) = \prod_{i=2}^{N} \frac{k_i}{s+q_i}$ is indeed the one given in Eq. (4). In order to do this, we write the function using partial fractions:

$$\prod_{i=2}^{N} \frac{k_i}{s+q_i} = k_p \sum_{i=2}^{N} \frac{B_i}{s+q_i} = k_p \frac{\sum_{i=2}^{N} B_i \left( \prod_{j \neq i} (s+q_j) \right)}{\prod_{i=2}^{N} (s+q_i)}.$$

We have used $k_p = \prod_i k_i$ here for notational convenience. We thus need to find the coefficients $B_i$ such that for all $s$

$$\sum_{i=2}^{N} B_i \left( \prod_{j \neq i} (s+q_j) \right) = 1.$$

This can be done by substituting $s = q_k$ for each $k$, which finally leads to

$$B_i = \frac{1}{\prod_{j \neq i} (q_j - q_i)}.$$

Using the known fact that the inverse Laplace transform of $1/(s+b)$ is $e^{-bt}$ and that the transform is linear, we find that

$$g(t) = \mathcal{L}^{-1} \left( k_p \sum_{i=1}^{N} \frac{B_i}{s+q_i} \right) = k_p \sum_{i=2}^{N} B_i e^{-q_i t} = k_p \sum_{i=2}^{N} \frac{1}{\prod_{j \neq i} (q_j - q_i)} e^{-q_i t} \tag{8}$$

which is indeed Eq. (4). Interestingly, only the parameters $q_i$ determine the shape of the distribution, and all the $k_i$ are grouped together into an amplification factor. Note that this distribution is not normalized:

$$\int_0^\infty g(t) \mathrm{d}t = \frac{\prod_{i=2}^{N} k_i}{\prod_{i=2}^{N} q_i}.$$

The normalized version of this distribution is called a hypoexponential distribution.

### 2 Calculation of the Hopf bifurcation lines

In this section, we provide the details of the calculation of the parameter values that lead to a Hopf bifurcation.

#### 2.1 Hopf bifurcation in the PI model

We first give the details of the calculation of the stability lines for the model

$$u' = cg(u_\theta) - u, \tag{9}$$

with $g(u) = 1/(1 + u^m)$. The steady state $\bar{u}$ satisfies $g(\bar{u}) = \bar{u}/c$. In order to study the stability of the steady state, we set $u = \bar{u} + \xi$, with $\xi$ a small perturbation, and substitute into Eq. (9):

$$(\bar{u} + \xi)' = cg(\bar{u} + \xi_\theta) - (\bar{u} + \xi).$$

Expanding the right hand side and keeping only terms linear in $\xi$ gives the linearized equation

$$\xi' = cg'(\bar{u})\xi_\theta - \xi. \tag{10}$$

In order to study the stability of the steady state, we check whether solutions of the linearized equation grow or decay in time. To this end, we substitute $\xi = e^{\lambda t}$, which gives the characteristic equation

$$\lambda + \alpha + \beta e^{-\lambda\theta} = 0, \tag{11}$$

with

$$\alpha = 1 \qquad \text{and} \qquad \beta = -cg'(\bar{u}).$$

Both these coefficients are positive. The steady state is linearly stable if all solutions of this equation have real part smaller than zero. Without delay, the only solution is $\lambda = -\alpha < 0$, which means that the solution is stable in the absense of delay. We look for a stability change as the delay increases. This can only happen through a Hopf bifurcation (MacDonald 2008), at which there are two purely imaginary solutions of the characteristic equation. Substituting $\lambda = i\omega$ and separating real and imaginary part leads to

$$\omega = \beta \sin(\omega\theta) \tag{12}$$

$$\alpha = -\beta \cos(\omega\theta), \tag{13}$$

which can be reworked to obtain

$$\omega = \sqrt{\beta^2 - \alpha^2} \tag{14}$$

$$\theta = \frac{1}{\omega} \arctan\left(-\frac{\omega}{\alpha}\right). \tag{15}$$

This gives an explicit method to compute the time delay that destabilizes a steady state. Given the parameters $m$ and $c$, we can compute the steady state $\bar{u}$, and from this $\alpha$ and $\beta$. Solving the equations above first for $\omega$ and then $\theta$ gives the frequency of the oscillation and the critical time delay. We can see that the equation for $\omega$ only has a real solution if $\beta > \alpha$. This corresponds to condition obtained from the geometrical picture in the main text: $\alpha/\beta < 1$.

To obtain the boundary lines in a two-parameter bifurcation diagram, we can fix $m$ or $c$, and perform the above recipe while varying the other parameter. This requires us to numerically solve for the steady state $\bar{u}$. There is a more elegant method that does not require a numerical solver to obtain $\bar{u}$. This method is also explained in the book by Erneux (2009). The idea is to parametrize the curves in parameter space by $\bar{u}$. For example, to obtain the line of instability in $(c, \theta)$-plane, we vary $\bar{u}$, and for each $\bar{u}$ we compute $c$ from the steady-state condition $c = \bar{u}/g(\bar{u})$. Since we choose $\bar{u}$, we can explicitly compute $\alpha, \beta$ and the time delay $\theta$. We thus have $c(\bar{u})$ and $\theta(\bar{u})$ which describe the curve in $(c, \theta)$-plane. Moreover, every $\bar{u} > 0$ corresponds to a single $c$, we we obtain the whole curve with this method. The same can be done for the diagrams in the $(m, \theta)$-plane.

##### 2.1.1 Asymptotics

The regime diagrams in the main text show that the stability boundary in the $(c, \theta)$-plane has a horizontal asymptote. This means that for large $c$, the critical time delay approaches a constant value. To compute this value, we consider the limit $c \to \infty$, and see how $\bar{u}$, $\alpha, \beta, \omega$ and $\theta$ scale in this limit.

We rewrite the steady state condition as

$$\bar{u}(1 + \bar{u}^m) = c. \tag{16}$$

For $c \to \infty$, a dominant balance argument shows that $\bar{u} \sim c^{1/(m+1)}$. We have that

$$\beta = -cg'(\bar{u}) = cm \frac{\bar{u}^{m-1}}{(1 + \bar{u}^m)^2}.$$

With the scaling $\bar{u} \sim c^{1/(m+1)}$, we find that $\beta \sim m$ for large $c$ and to leading order, implying $\omega \sim \sqrt{m^2 - 1}$. We substitute this in the expression for the critical delay (Eq. (15)):

$$\theta \to \frac{\arctan\left(-\sqrt{m^2 - 1}\right)}{\sqrt{m^2 - 1}} \quad \text{for} \quad c \to \infty. \tag{17}$$

Here we have to keep in mind to use values of arctan between 0 and $\pi$.

The boundary in the $(c, \theta)$-plane also has a vertical asymptote, corresponding to the limit $\theta \to \infty$. We can compute the corresponding value of $c$. Eq. (15) implies that if $\theta \to \infty$, $\omega \to 0$, or $\alpha \to \beta$ (from Eq. (14)). So, we require

$$1 = -cg'(\bar{u}).$$

Combined with the steady state condition $cg(\bar{u}) = \bar{u}$ and some algebra, this leads to

$$c = m(m - 1)^{-1-1/m}, . \tag{18}$$

This is the lowest value of $c$ for which there exists a destabilizing time delay, and thus the lowest value of $c$ for which oscillations are possible, given $m$. Fig. S1 shows these asymptotes.

We can also compute an approximation for the destabilizing time delay for large $m$. For this computation, we use the following approximation (inspired by Woller, Gonze, and Erneux (2013)):

$$(1 + \frac{x}{m})^m = e^x + \mathcal{O}(1/m) \quad \text{for } m \to \infty. \tag{19}$$

This can be shown as follows. Take the logarithm of the expression $(1 + x/m)^m$ and expand in a Taylor series (considering $x$ constant and $m$ large):

$$\begin{aligned} m\ln(1+x/m) &= m\left(\frac{x}{m} + \mathcal{O}\left(\frac{x}{m}\right)^2\right) \\ &= x + \mathcal{O}\left(\frac{x^2}{m}\right) \end{aligned} \tag{20}$$

This means (now also expanding the exponential function around zero)

$$(1+\frac{x}{m})^m = e^{x+\mathcal{O}(x^2/m)} = e^x(1 + \mathcal{O}(x^2/m) + \frac{1}{2}\mathcal{O}(x^4/m^2) + \cdots) = e^x + \mathcal{O}(1/m) \tag{21}$$

for a fixed $x$ and as $m \to \infty$.

To compute an approximation for the critical time delay for large $m$, we assume that $c > 1$. In this case, the steady state $\bar{u}$ approaches 1 as $m \to \infty$. First, we seek an expansion of $\bar{u}$ for large $m$. We assume it takes the form $\bar{u} = 1 + u_1 m^{-1} + \mathcal{O}(m^{-2})$. The equation for the steady state reads (keeping only terms up to order $1/m$)

$$1 + u_1 m^{-1} = cg(1 + u_1 m^{-1}) = c\frac{1}{1 + (1 + u_1 m^{-1})^m}.$$

Using the formula above, this reduces to

$$1 + u_1 m^{-1} = c\frac{1}{1 + e^{u_1} + \mathcal{O}(1/m)} = c\frac{1}{1 + e^{u_1}} + \mathcal{O}(1/m).$$

Comparing the leading terms leads to

$$u_1 = \ln(c-1) \tag{22}$$

such that $\bar{u} = 1 + \ln(c-1)\frac{1}{m} + \mathcal{O}(m^{-2})$ for $m \to \infty$. Using this, we can compute the leading terms of $\alpha$ and $\beta$ and then $\omega$ and $\theta$. For this model, $\alpha = 1$ and

$$\beta = -cg'(\bar{u}) = cm\frac{\bar{u}^{m-1}}{(1+\bar{u}^m)^2}.$$

Substituting the asymptotic formula for $\bar{u}$ yields that, to leading order,

$$\beta \sim m\frac{c-1}{c}.$$

From this, we find

$$\omega = \sqrt{\beta^2 - \alpha^2} \sim m\frac{c-1}{c}$$

and

$$\theta = \frac{1}{\omega}\arctan(-\omega/\alpha) \sim \frac{c}{c-1}\frac{\pi}{2}\frac{1}{m}, \tag{23}$$

as $m \to \infty$. Here we used that arctan goes to $\pi/2$, taking into account that we need values in $[0, \pi]$. Note that for large $c$, this expression goes to $\pi/(2m)$. This is the same value as the limit of Eq. (17) for large $m$. The approximations for both $m$ and $c$ large are thus consistent.

We summarize the different approximations for the PI model:

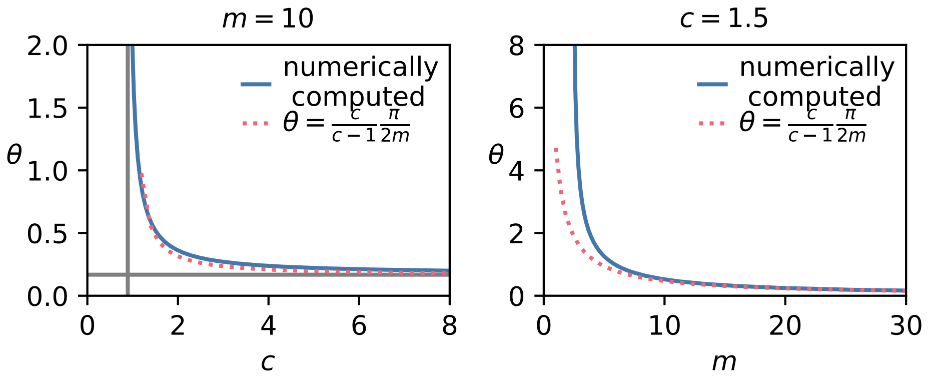


Figure S1: Stability boundaries for the PI model with asymptotic approximations. Left: stability boundary in the $(c, \theta)$-plane. The vertical asymptote is $c = m(m-1)^{-1-1/m}$ and the horizontal asymptote is $\theta = \frac{\arctan\left(-\sqrt{m^2-1}\right)}{\sqrt{m^2-1}}$. The dotted line shows the approximation for large $m$ given by $\theta = \frac{c}{c-1}\frac{\pi}{2m}$. Right: stability boundary in the $(m, \theta)$-plane with approximation for large $m$. The dashed and solid lines coincide as $m \to \infty$.

- $\theta_c \to \frac{\arctan\left(-\sqrt{m^2-1}\right)}{\sqrt{m^2-1}}$ for fixed $m$ and $c \to \infty$
- $\theta_c \to \frac{c}{c-1}\frac{\pi}{2}\frac{1}{m}$ for fixed $c > 1$ and $m \to \infty$
- For fixed $m$, the critical time delay $\theta_c$ goes to infinity as $c \xrightarrow{>} m(m-1)^{-1-1/m}$

Fig. S1 summarizes these asymptotic results.

#### 2.2 Hopf bifurcation in the DA model

The approach is identical to the one for the PI model. This calculation can also be found in our previous paper (Rombouts, Vandervelde, and Gelens 2018). For this model, the governing equation is

$$u' = c - f(u_\theta)u, \qquad f(u) = \frac{u^m}{1+u^m}. \tag{24}$$

The equation, linearized around the steady state $\bar{u}$, reads

$$\xi' = -f(\bar{u})\xi - \bar{u}f'(\bar{u})\xi_\theta. \tag{25}$$

The characteristic equation is thus

$$\lambda + \alpha + \beta e^{-\lambda\theta} = 0 \tag{26}$$

with $\alpha = f(\bar{u})$ and $\beta = \bar{u}f'(\bar{u})$. The way of determining instability curves is the same as for the PI model. In particular, Eqs. (14) and (15) are exactly the same. As before, we can parametrize the curves in two-parameter bifurcation diagrams by $\bar{u}$.

##### 2.2.1 Asymptotics

Contrary to the PI model, the boundary curve in the $(c, \theta)$-plane does not have horizontal asymptotes, but rather two vertical asymptotes.

In order to compute the location of the vertical asymptotes of this curve in the $(c, \theta)$-plane, we look at the limit $\theta \to \infty$. From Eq. (15) this implies $\omega \to 0$, or $\alpha \to \beta$ (from Eq. (14)). This implies

$$f(\bar{u}) = \bar{u} f'(\bar{u}), \tag{27}$$

which is satisfied for $\bar{u} = 0$, corresponding to $c = 0$, or for

$$\bar{u}^m = m - 1.$$

The latter equation implies

$$c = \frac{(m-1)^{1+1/m}}{m}, \tag{28}$$

which is the location of the right asymptote. As is expected due to the equivalence result between the two models, this value is the inverse of the value obtained for the vertical asymptote for the PI model (Eq. (18)).

Eqs (18) and Eq. (28) also describe the boundary in the $(c, m)$ plane above which a destabilizing time delay exists (Fig. 3(l) in the main text).

Interestingly, when considering $c$ as function of $m$, Eq. (28) is non-monotonic. Rather, it has a maximal value at $m \approx 10.19$. For this value of $m$, the range of $c$-values for which oscillations are possible is maximal.

As for the PI model, we can also obtain an approximate expression for the critical delay for large values of the Hill exponent $m$. As before, we first approximate the value of the steady state for large $m$. We take $c < 1$ and assume an expression of the form $\bar{u} = 1 + u_1 m^{-1}$. The steady state equation is

$$c = \bar{u} f(\bar{u}) = (1 + u_1 m^{-1}) \frac{(1 + \frac{u_1}{m})^m}{1 + (1 + \frac{u_1}{m})^m} = \frac{1}{1 + e^{-u_1}} + \mathcal{O}(m^{-1}),$$

where we used Eq. (19). Balancing the leading order terms gives $u_1 = \ln(c/(1-c))$ and thus

$$\bar{u} = 1 + \ln\left(\frac{c}{1-c}\right)\frac{1}{m} + \mathcal{O}(m^{-2}).$$

Substituting this expression into $\alpha = f(\bar{u})$ and $\beta = \bar{u} f'(\bar{u})$, we find that to leading order

$$\alpha \sim c \tag{29}$$
$$\beta \sim mc(1-c). \tag{30}$$

This gives

$$\omega = \sqrt{\beta^2 - \alpha^2} \sim mc(1-c)$$

and

$$\theta \sim \frac{1}{c(1-c)} \frac{\pi}{2} \frac{1}{m} \tag{31}$$

using the same reasoning for the arctangent as before.

Fig. S2 shows the asymptotic approximations.

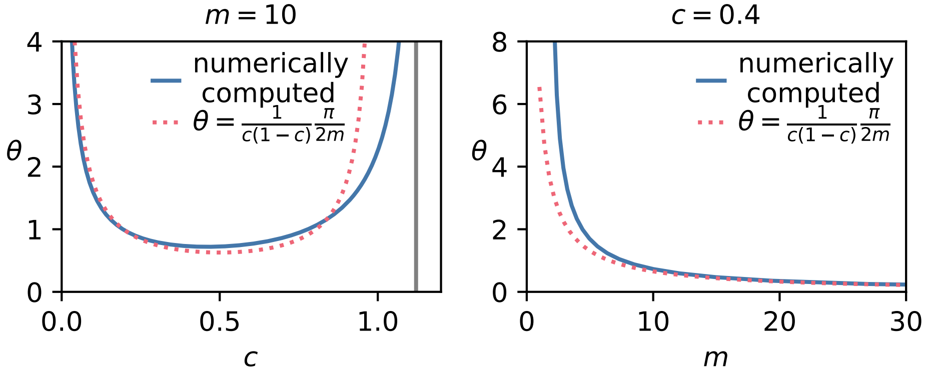


Figure S2: Stability boundaries for the DA model with asymptotic approximations. Left: stability boundary in the $(c, \theta)$-plane. The vertical asymptote is $c = \frac{(m-1)^{1+1/m}}{m}$. The dotted line shows the approximation for large $m$ given by $\theta = \frac{1}{c(1-c)}\frac{\pi}{2m}$. Right: stability boundary in the $(m, \theta)$-plane with approximation for large $m$. The dashed and solid lines coincide as $m \to \infty$.

#### 2.3 Hopf bifurcation for Gamma-distributed delay

We consider now the models with a distributed delay

$$\begin{aligned} u' &= cg(u_\theta) - u \\ u' &= c - f(u_\theta)u \end{aligned} \tag{32}$$

with

$$u_\theta = \int_0^\infty g_a^N(s)u(t-s)\mathrm{d}s.$$

Here $g_a^N$ is a Gamma distribution with density function

$$g_a^N(s) = \frac{a^N}{(N-1)!}s^{N-1}e^{-as}.$$

In the main text we consider the gamma distribution with fixed mean. If the mean is fixed at $\bar{\theta}$, the parameter $a$ is equal to $N/\bar{\theta}$ and increasing $N$ makes the distribution more sharply peaked, with the $N \to \infty$ limit being a delta distribution.

The linearization of Eqs. (32) around the steady state $\bar{u}$ is as before. The resulting linear equation for both models is

$$\xi' = -\alpha\xi - \beta\xi_\theta \tag{33}$$

where $\xi_\theta$ is now also given by the convolution with $g_a^N$ and the values of $\alpha$ and $\beta$ are the same as for the corresponding models with a discrete delay.

Now, the exponential ansatz $\xi = e^{\lambda t}$ yields

$$\lambda = -\alpha - \beta\mathcal{L}(g_a^N)(\lambda), \tag{34}$$

where

$$\mathcal{L}(g_a^N) = \int_0^\infty e^{-\lambda s} g_a^N(s)\mathrm{d}s$$

is the Laplace transform of $g_a^N$ and $\alpha$ and $\beta$ are the same as before.

For a Gamma distribution the Laplace transform is known:

$$\mathcal{L}(g_a^N)(\lambda) = \frac{a^N}{(\lambda + a)^N} = (1 + (\lambda/a))^{-N}.$$

With $a = N/\bar{\theta}$, the characteristic equation (33) can be written as

$$\lambda + \alpha + \beta \left(1 + \frac{\lambda\bar{\theta}}{N}\right)^{-N} = 0 \tag{35}$$

With the Hopf ansatz $\lambda = i\omega$ we can write the equation as

$$-\frac{\alpha}{\beta} - i\frac{\omega}{\beta} = (1 + \frac{i\omega\bar{\theta}}{N})^{-N}. \tag{36}$$

As described in the main text, the right hand side describes a spiral in the complex plane, parameterized by $\omega\bar{\theta}$. The left hand side describes a vertical line in the lower complex plane. Fig. S3 shows some of these spirals for different values of $N$. For large $N$, the spirals approximate the circle in the bottom half-plane.

For further analysis, it is convenient to define

$$1 + \frac{i\omega\bar{\theta}}{N} = Re^{i\gamma}$$

with

$$R^2 = 1 + \left(\frac{\omega\bar{\theta}}{N}\right)^2 \qquad \text{and} \qquad \tan\gamma = \frac{\omega\bar{\theta}}{N}.$$

We also have $R^2 = 1 + \tan^2\gamma = \frac{1}{\cos^2\gamma}$ Note that since $\omega, \bar{\theta}$ and $N$ are positive, we consider $\gamma \in [0, \pi/2]$. This entails

$$R\cos\gamma = 1.$$

The spiral can now be parametrized by $\gamma$:

$$(1 + \frac{i\omega\bar{\theta}}{N})^{-N} = \cos^N(\gamma)e^{-iN\gamma} = \cos^N(\gamma)(\cos(N\gamma) - i\sin(N\gamma))$$

The point at which the Hopf bifurcation occurs can be found by finding the values of $\omega$ and $\gamma$ such that the vertical line and the spiral intersect:

$$-\frac{\alpha}{\beta} - i\frac{\omega}{\beta} = \cos^N(\gamma)(\cos(N\gamma) - i\sin(N\gamma)).$$

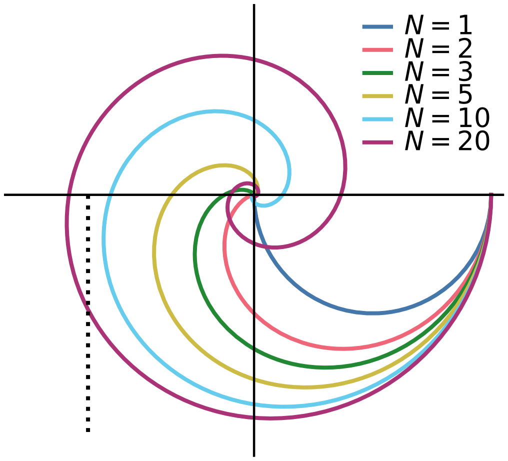


Figure S3: The right hand side of Eq. (36) describes a spiral in the complex plane for varying $\omega\bar{\theta}$. The spirals all start at $(1, 0)$. A Hopf bifurcation is possible if the spiral intersects with the vertical line with real part $-\alpha/\beta$.

Separating real part and imaginary part yields

$$-\alpha = \beta \cos^N(\gamma)\cos(N\gamma) \tag{37}$$

$$\omega = \beta \cos^N(\gamma)\sin(N\gamma) \tag{38}$$

These equations need to be solved for $\omega$ and $\gamma$. Then $\bar{\theta}$ can be computed as $\bar{\theta} = N(\tan\gamma)/\omega$. In this way, the bifurcation curves can be obtained.

The first of the equations can be rewritten by setting $y = \cos\gamma$. The equation then reads

$$-\alpha = \beta y^N T_N(y), \tag{39}$$

where $T_N$ is the $N$-th Chebyshev polynomial of the first kind.

One procedure to compute the bifurcation curve in the $(c, \bar{\theta})$-plane, parametrized by $\bar{u}$, is as follows.

1. Pick $\bar{u}$ and compute associated $\alpha$ and $\beta$.
2. Solve Eq. (39) numerically for $y$, requiring $y \in [\arccos(\pi/(2N)), \arccos(\pi/N]$. This interval is the one that corresponds to intersections in the lower left quadrant where $N\gamma \in [\pi/2, \pi]$.
3. Compute $\omega$ from the identity $\omega^2 = \beta^2 y^{2N} - \alpha^2$.
4. Compute $\bar{\theta}$ from
$$\bar{\theta} = \frac{N}{\omega}\tan\gamma = \frac{N}{\omega}\sqrt{y^{-2} - 1}$$
5. The corresponding value of $c$ can be found directly from the steady state conditions, $cg(\bar{u}) = \bar{u}$ for the PI model and $c = \bar{u}f(\bar{u})$ for the DA model.

The same procedure can be used to find the bifurcation curve in the $(m, \bar{\theta})$-plane. A caveat is that there can be zero, one or two solutions of the equation $-\alpha = \beta y^N T_N(y)$. All the relevant solutions have to be found to determine the complete bifurcation curve. See also the discussion of the number of intersections in Section 2.4.

An alternative method, which circumvents the problem of numerically finding the solutions of the equation involving the Chebyshev polynomials, is to parametrize the instability curve by $\gamma$. Each value of $\gamma$ correspond to a single point on the spiral, which has an angle of $-N\gamma$ with the horizontal axis. We only look at points in the lower left quadrant which have $-\pi \leq -N\gamma \leq -\pi/2$, or $\gamma$ between $\pi/(2N)$ and $\pi/N$.

The procedure for computing the bifurcation line in the $(c, \bar{\theta})$ plane then goes as follows.

1. Fix $\gamma \in [\pi/(2N), \pi/N]$.
2. Compute the real part of the intersection point as $x = \cos^N(\gamma)\cos(N\gamma)$ and the imaginary part as $y = -\cos^N(\gamma)\sin(N\gamma)$.
3. The real part of the vertical line is $-\alpha/\beta$, therefore we find $\alpha/\beta = -\cos^N(\gamma)\cos(N\gamma)$. For the PI model, $\alpha/\beta = \frac{1}{m}(1 + \bar{u}^{-m})$, which can be solved (for fixed $m$) for $\bar{u}$. Then $c$ can be determined from $cg(\bar{u}) - \bar{u} = 0$. For the DA model, $\alpha/\beta = \frac{1}{m}(1 + \bar{u}^m)$ which can be solved for $\bar{u}$, and $c$ can be found from $c - \bar{u}f(\bar{u}) = 0$.

4. It is now possible to compute $\beta$. For the PI model $\beta = -cg'(\bar{u})$ and for the DA model $\beta = \bar{u}f'(\bar{u})$. By requiring imaginary parts of vertical line and spiral to be equal, find $\omega = \beta \cos^N(\gamma)\sin(N\gamma)$.

5. Given $\omega$ and $\gamma$, we can now compute $\bar{\theta}$ as $\bar{\theta} = N \tan\gamma/\omega$.

The benefit of this procedure is that we do not need a numerical solver for the roots of an equation, and that we find the whole bifurcation curve. It works for the curve in the $(c, \bar{\theta})$ plane because of step 3 above: given $\alpha/\beta$, the value of $\bar{u}$ can be found if $m$ is fixed, and $c$ can be computed afterwards. To compute the bifurcation curve in the $(m, \theta)$-plane step 3 would be to solve (example for PI model) the system

$$\frac{\alpha}{\beta} = \frac{1}{m}(1 + \bar{u}^{-m})$$
$$c = \bar{u}(1 + \bar{u}^m)$$

for $\bar{u}$ and $m$ simultaneously.

To determine the region in the $(c, m)$-plane where oscillations exist, we solve $-\alpha/\beta = Q(N)$ where $Q(N)$ is the minimal real part of the spiral, which depends on $N$. The real part of the spiral is given by $\cos^N \gamma \cos(N\gamma)$ which is minimal for $\gamma = \frac{\pi}{N+1}$. So we find that

$$Q(N) = \cos^N \frac{\pi}{N+1} \cos \frac{N\pi}{N+1} = -\cos^{N+1} \frac{\pi}{N+1}.$$

To obtain the boundaries in the $(c, m)$-plane, we need to solve $\alpha/\beta = Q(N)$ together with the steady state condition. This leads to

$$c = mQ(N)(mQ(N) - 1)^{-1-1/m} \tag{40}$$

for the PI model and

$$c = \frac{1}{mQ(N)}(mQ(N) - 1)^{1+1/m} \tag{41}$$

for the DA model. These curves are shown in the main text, Fig. 4(c).

#### 2.4 Hopf bifurcation for the system with double regulation

In this section we describe the Hopf bifurcation for the system with regulation both in the production and inhibition terms. If the threshold and exponent of the Hill functions, as well as the time delay, are equal, the scaled system is

$$u' = cg(u_\theta) - f(u_\theta)u. \tag{42}$$

The steady state satisfies $cg(\bar{u}) = f(\bar{u})\bar{u}$ which means $c = \bar{u}^{m+1}$. For this system, the steady state can thus be calculated analytically. The linear stability goes as always and we have the characteristic equation

$$\lambda + \alpha + \beta e^{-\lambda\theta} = 0,$$

with

$$\alpha = f(\bar{u}) \qquad \text{and} \qquad \beta = -cg'(\bar{u}) + f'(\bar{u})\bar{u}.$$

We can rewrite $\beta$ as $m\bar{u}^m/(1+\bar{u}^m)$ and therefore $\frac{\alpha}{\beta} = \frac{1}{m}$. Since $\alpha/\beta$ determines the real part of the intersection point, in the graphical description of the Hopf bifurcation, we see that the existence of a time delay that destabilizes the steady state does not depend on $c$, only on $m$. The value of $\theta$ that destabilizes the steady state does, however, depend on $c$ and can be computed as before.

In the system with Gamma-distributed delay, we can compute for which values of $m$ the vertical line has zero, one or two intersections with the spiral. The left panel of Fig. S4 illustrates this for $N = 3$. In this figure, point $P1$ with coordinates $(x_1, y_1)$ is the point with minimal real part of the spiral and $P2$ with coordinates $(x_2, y_2)$ is the point where the spiral first crosses the negative horizontal axis. The vertical line has real part $-\alpha/\beta$. It is clear that no intersection exists if $-\alpha/\beta < x_1$, that two intersections exist if $x_1 < -\alpha/\beta < x_2$ and that a single intersection exists if $-\alpha/\beta > x_2$. Note: for larger $N$ (e.g. $N = 20$ in Fig. S3) more intersections are possible. However only the first two correspond to stability switches, see MacDonald (2008).

The spiral, parameterized by $\gamma$, has real part $x = \cos^N \gamma \cos(N\gamma)$ and imaginary part $y = -\cos^N \gamma \sin(N\gamma)$ (see Section 2.3). As discussed in the previous section, the minimum real part is given by $Q(N) = -\cos^{N+1} \frac{\pi}{N+1}$. The point $P_2$ has $\gamma = \pi/N$. This means that

$$\begin{aligned} x_1 &= -\cos^{N+1} \frac{\pi}{N+1} & y_1 &= -\cos^N \frac{\pi}{N+1} \sin \frac{N\pi}{N+1} \\ x_2 &= -\cos^N \frac{\pi}{N} & y_2 &= 0 \end{aligned} \tag{43}$$

By comparing $x_1$ and $x_2$ to $\alpha/\beta = 1/m$, we find the conditions on $m$ and $N$ for two intersections to exist: $x_1 < -1/m < x_2$ or

$$-\cos^{N+1} \frac{\pi}{N+1} < -1/m < -\cos^N \frac{\pi}{N}, \tag{44}$$

which can be rewritten as

$$\cos^{-(N+1)} \frac{\pi}{N+1} < m < \cos^{-N} \frac{\pi}{N}. \tag{45}$$

These bounds are also illustrated in Fig. S4.

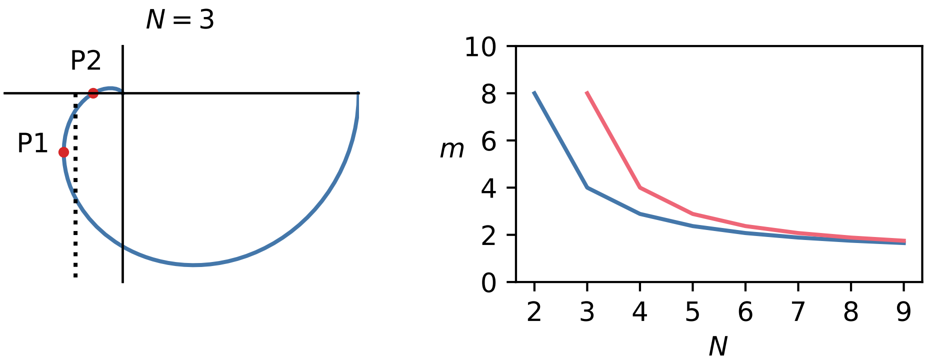


Figure S4: Left: illustration of the points whose coordinates are given in Eq. (43), to determine how many intersection points the vertical line has. Right: values of $m$ for which oscillations are possible as function of $N$ in the double regulation system, illustrating Eq. (45). For values of $m$ above the blue line, there is a delay that destabilizes the steady state. For values of $m$ in between the blue and red lines, there exists a delay, larger than the destabilizing delay, that restabilizes the steady state.

### 3 Calculations for the limit of large Hill exponent

In the limit of large Hill exponents, the Hill function can be approximated by a step, or Heaviside function. This makes the system analytically solvable (Mackey 1997)(Erneux 2009, pp. 56-59). The main idea is to solve the simplified equation, where the step function is either zero or one, on different parts, and afterwards glue these parts together by requiring continuity of the solution.

We define the interval $[0, \bar{t}]$ as the interval on which the solution is decreasing, and $[\bar{t}, P]$ the interval on which the solution is increasing. The maximum and minimum of the oscillation are $u_{\max}$ and $u_{\min}$. Fig. S5 shows the situation for the PI and DA model. The calculation to be done is to use the delay differential equation to obtain equations for the four values $\bar{t}, P, u_{\max}$ and $u_{\min}$.

#### 3.1 PI model

The equation for large $m$ is

$$u' = c(1 - H(u_\theta - 1)) - u, \tag{46}$$

where $H$ is the Heaviside function. On the first region we have $u' = -u$ with solution $u(t) = u_{\max}e^{-t}$, on the second part the equation is $u' = c - u$ with solution $u(t) = c + (u_{\min} - c)e^{-(t-\bar{t})}$. The set of equations for $u_{\max}, u_{\min}, \bar{t}$ and $P$ reads

$$u_{\min} = u_{\max}e^{-\bar{t}} \tag{47}$$

$$u_{\max} = c + (u_{\min} - c)e^{-(P-\bar{t})} \tag{48}$$

$$u_{\max}e^{-(\bar{t}-\theta)} = 1 \tag{49}$$

$$c + (u_{\min} - c)e^{-(P-\theta-\bar{t})} = 1. \tag{50}$$

We can solve this and find

$$u_{\min} = e^{-\theta} \tag{51}$$

$$u_{\max} = c + (1 - c)e^{-\theta} \tag{52}$$

$$\bar{t} = \ln\left(\frac{u_{\max}}{u_{\min}}\right) \tag{53}$$

$$P = \bar{t} + \ln\left(\frac{u_{\min} - c}{u_{\max} - c}\right). \tag{54}$$

This gives us, for the amplitude $A = u_{\max} - u_{\min}$ and period:

$$A = c(1 - e^{-\theta}) \tag{55}$$

$$P = 2\theta + \ln(c + (1 - c)e^{-\theta}) + \ln\left(\frac{e^{-\theta} - c}{1 - c}\right) \tag{56}$$

#### 3.2 DA model

The equation for large $m$ becomes

$$u' = c - H(u_\theta - 1)u, \tag{57}$$

where $H$ is the Heaviside function. On the decreasing part, the equation for $u$ is

$$u' = c - u,$$

with solution

$$u(t) = c + (u_{\max} - c)e^{-t}.$$

on the increasing part $u' = c$ with solution

$$u(t) = u_{\min} + c(t - \bar{t}).$$

From this, we derive the following set of equations for $u_{\max}, u_{\min}, \bar{t}$ and $P$:

$$u_{\min} = c + (u_{\max} - c)e^{-\bar{t}} \tag{58}$$

$$u_{\max} = u_{\min} + c(P - \bar{t}) \tag{59}$$

$$c + (u_{\max} - c)e^{-(\bar{t}-\theta)} = 1 \tag{60}$$

$$u_{\min} + c(P - \theta - \bar{t}) = 1. \tag{61}$$

The solution is

$$u_{\min} = c + (1 - c)e^{-\theta} \tag{62}$$

$$u_{\max} = 1 + c\theta \tag{63}$$

$$\bar{t} = \ln\left(\frac{u_{\max} - c}{u_{\min} - c}\right) \tag{64}$$

$$P = \bar{t} + \frac{u_{\max} - u_{\min}}{c}. \tag{65}$$

From this we find expressions for the amplitude $A = u_{\max} - u_{\min}$ and period $P$ as function of the parameters:

$$A = (1 - c)(1 - e^{-\theta}) + c\theta \tag{66}$$

$$P = 2\theta + \ln\left(1 + \frac{c}{1-c}\theta\right) + \frac{1-c}{c}(1 - e^{-\theta}) \tag{67}$$

### 4 Analysis of the system with enzymatic degradation

#### 4.1 Hopf bifurcation

The PI model with enzymatic degradation and small basal degradation reads

$$\frac{dX}{dt} = k_p \frac{K^m}{K^m + X_\tau^m} - k_d \frac{X}{1 + \gamma X} - k_b X. \tag{68}$$

For the DA model we have

$$\frac{dX}{dt} = k_p - k_d \frac{X_\tau^m}{K^m + X_\tau^m} \frac{X}{1 + \gamma X} - k_b X. \tag{69}$$

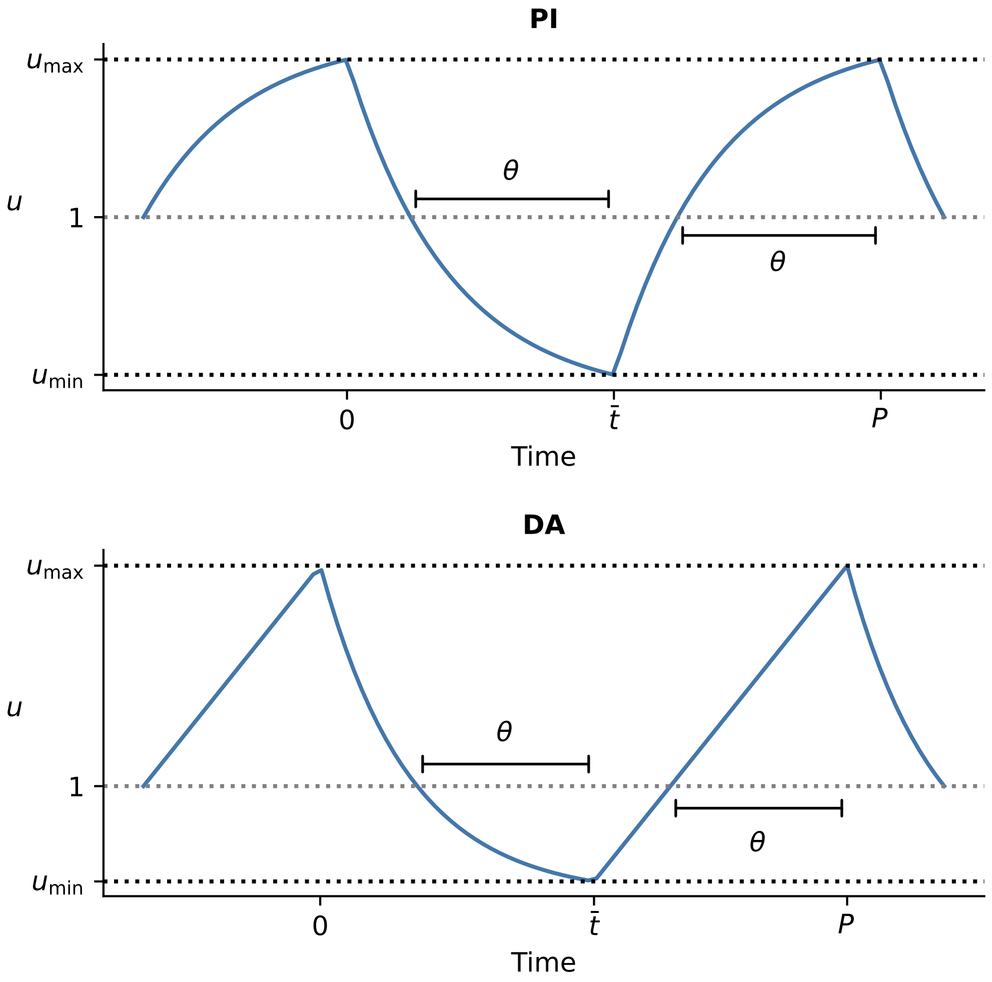


Figure S5: Analytical solution for the limit cycle. Top: for the PI model, bottom: for the DA model.

There are two degradation terms, the saturating one and the basal one. Only the saturating degradation term is regulated. Performing the scaling $u = X/K, s = k_d t$ gives

$$u' = cg(u_\theta) - h(u) - du \tag{PI}$$
$$u' = c - f(u_\theta)h(u) - du, \tag{DA}$$

with $c = k_p/(k_d K)$, $\theta = k_d\tau$, $d = k_b/k_d$ and

$$h(u) = \frac{u}{1 + \kappa u} \text{ with } \kappa = K\gamma. \tag{70}$$

For the PI model, the steady state satisfies

$$cg(u) = h(u) + du$$

and for the DA model

$$f(u)h(u) = c - du.$$

If $d > 0$, there is a unique positive steady state. If $d > 0$, it is possible that the DA model does not admit a steady state if $c$ is too large.

A linear stability analysis leads to the characteristic equation

$$\lambda + \alpha + \beta e^{-\lambda\theta} = 0$$

with

$$\alpha = h'(\bar{u}) + d \qquad \beta = -cg'(\bar{u}) \qquad \text{(PI model)}$$
$$\alpha = f(\bar{u})h'(\bar{u}) + d \qquad \beta = f'(\bar{u})h(\bar{u}). \qquad \text{(DA model)}$$

The Hopf bifurcation analysis is completely analogous to the basic model. Stability boundaries in parameter planes can again be computed when they are parametrized by $\bar{u}$. An example for the PI model: to compute the stability region in the $(m, \theta)$-plane for fixed $c$, use $\bar{u}$ as a varying parameter. For each $\bar{u}$, compute $\bar{g}$ as $(h(\bar{u}) + d\bar{u})/c$ (all the parameters here are known). Then $m$ can be computed from $\frac{1}{1+\bar{u}^m} = \bar{g}$:

$$m = \ln(1/\bar{g} - 1)/\ln\bar{u}.$$

The values of $\alpha$ and $\beta$ can now be computed, and from this $\omega$ and $\theta$.

#### 4.2 The oscillatory region in the $(c, m)$-plane

In the main text, we have shown regions in the $(c, m)$-plane for which oscillations are possible. This means, for these values of $c$ and $m$ there exists a $\theta$ that corresponds to the Hopf bifurcation. Form the geometrical interpretation of the Hopf bifurcation, we know that — for the discrete delay models — this is only possible if $\alpha/\beta < 1$. The boundary of this region in the $(c, m)$-plane thus corresponds to $\alpha = \beta$.

For the PI model, the values of $c$ and $m$ on this boundary satisfy the system of equations given by the steady-state condition and $\alpha = \beta$. Written out, this is

$$h(u) + du = cg(u) \tag{71}$$

$$h'(u) + d = -cg'(u), \tag{72}$$

where we used $u$ instead of $\bar{u}$ for the steady state. These equations have three unknowns: $c, m$ and $u$, We can eliminate $c$, which leads to

$$-\frac{g'(u)}{g(u)} = \frac{h'(u) + d}{h(u) + du}. \tag{73}$$

We can rewrite this to

$$\frac{mu^{m-1}}{1 + u^m} = \frac{1}{u\zeta} \frac{1 + d\zeta^2}{1 + d\zeta} \tag{74}$$

where $\zeta = 1 + \kappa u$ for notational convenience. We then write this equality as

$$1 + u^m - mu^m\zeta + d\zeta^2(1 + (1 - m)u^m) = 0 \tag{75}$$

or

$$F(u, m) = 0,$$

with $F$ the expression on the left-hand side.

The boundary curve is now computed by tracking the $u, m$ such that $F(u, m) = 0$ using pseudo-arclength continuation (Kuznetsov 2004). For each $u$ and $m$ the value of $c$ can be computed as $c = (h(u) + du)/g(u)$.

A very similar approach is used for the DA model. The values of $m, c$ and $u$ now have to satisfy the system of equations

$$c = f(u)h(u) + du \tag{76}$$

$$f(u)h'(u) + d = f'(u)h(u) \tag{77}$$

The second equation can be written as

$$F(u, m) = u^m(1 + u^m - m\zeta) + d\zeta^2(1 + u^m)^2 = 0 \tag{78}$$

with $\zeta = 1 + \kappa u$. This curve can be tracked in the $(u, m)$-plane, and corresponding values of $c$ can be computed from the steady-state condition.

Figs. S6 and S7 show these curves for different values of $d$ and $\kappa$.

#### 4.3 Parameter set for which there is a stable steady state and a limit cycle

As an illustration of the more complex behavior possible in the models with enzymatic degradation, we describe here an example of bistability between steady state and limit cycle that we have observed by numerical simulation. For the set of parameters in Table S1, in the PI model, the

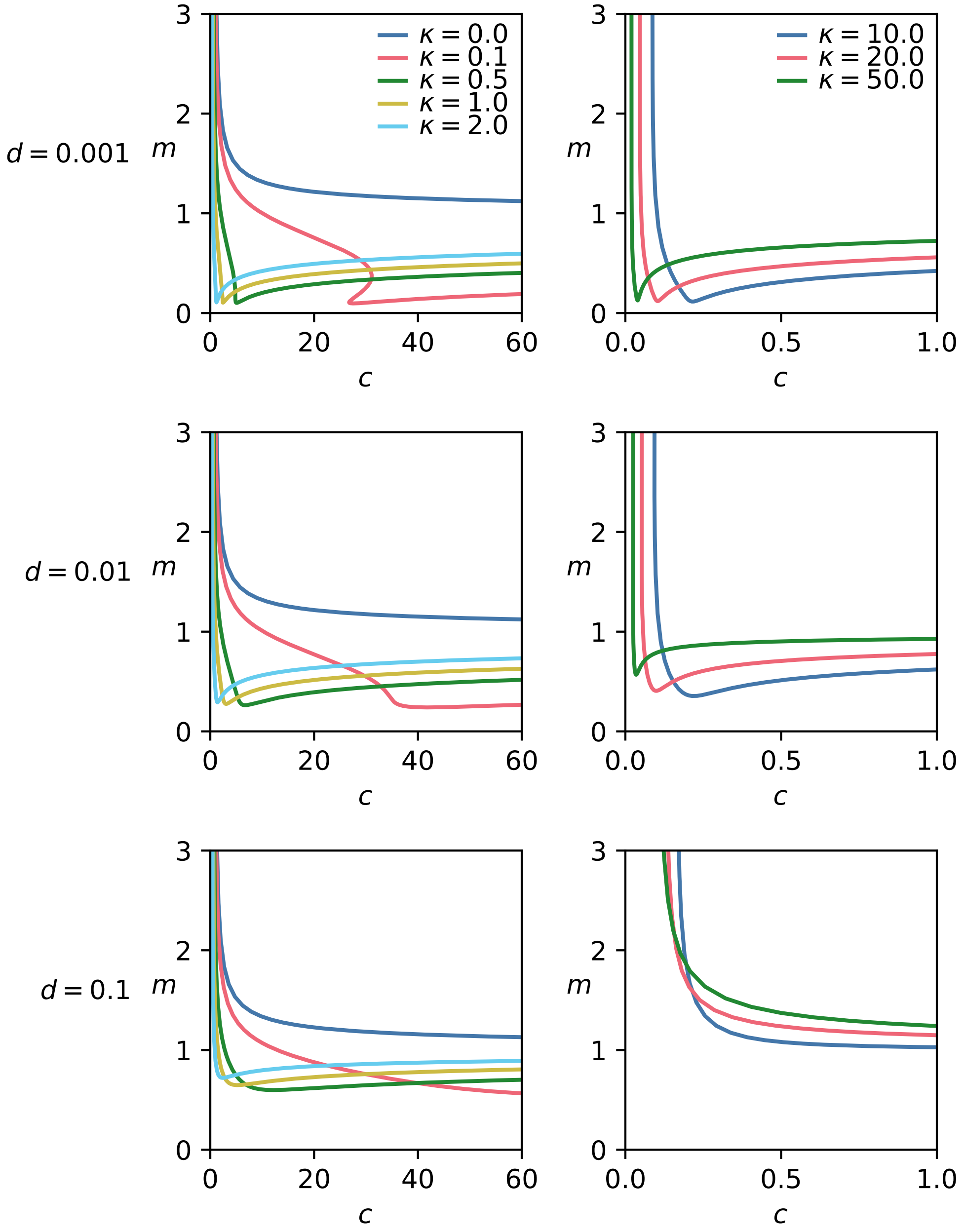


Figure S6: Stability boundaries in the $(c, m)$-plane for the PI model. Different rows correspond to different values of $d$, left is small values of $\kappa$, right is larger values of $\kappa$. Note the different axis scalings. The legends in the top panels hold for the whole column.

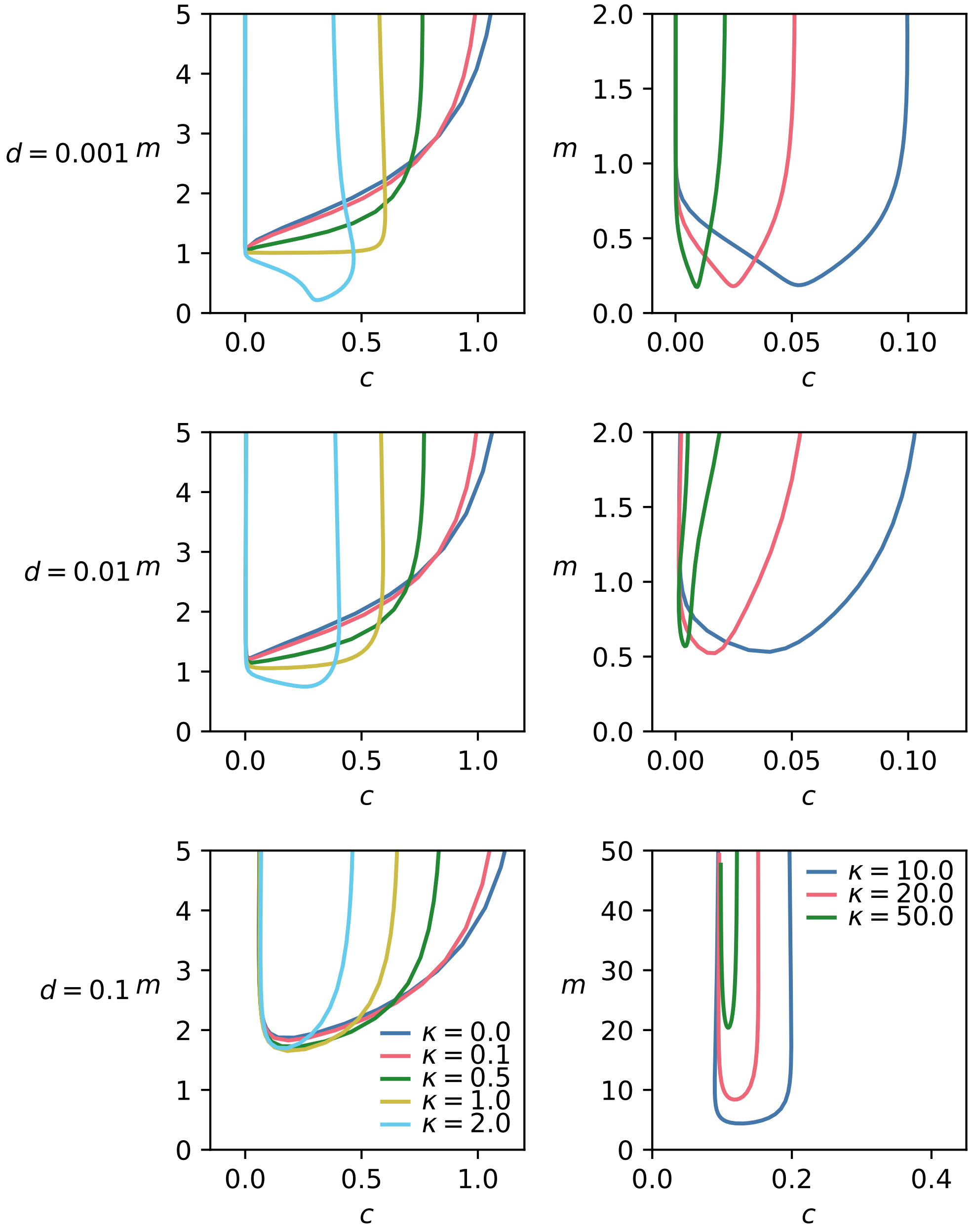


Figure S7: Stability boundaries in the $(c, m)$-plane for the DA model. Different rows correspond to different values of $d$, left is small values of $\kappa$, right is larger values of $\kappa$. Note the different axis scalings. The legends in the bottom panels hold for the whole column.

| Parameter | Value |
|---|---|
| $m$ | 0.45 |
| $c$ | 30 |
| $\kappa$ | 0.1 |
| $d$ | 0.001 |
| $\theta$ | 200 |

Table S1: Example of parameter values that give bistability between a steady state and an oscillation in the PI model with enzymatic degradation.

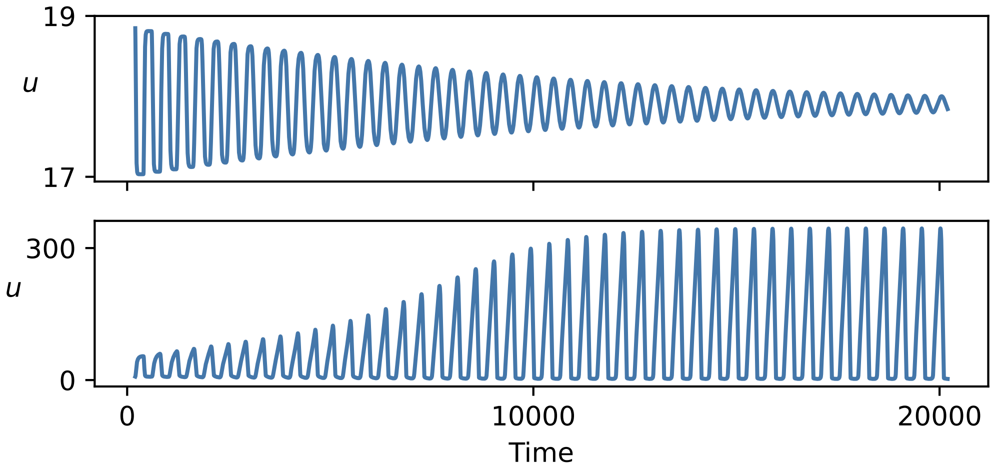


Figure S8: Bistability between steady state and limit cycle for specific set of parameter values (given in Table S1. Top: initial condition close to the steady state, bottom: initial condition farther away from the steady state. Note the difference in $y$-axis scale.

steady state is stable. The value of the steady state $\bar{u} \approx 17.9$ and the value of $\alpha/\beta \approx 1.019$. This is larger than one, which means a Hopf bifurcation is not possible. This steady state is thus linearly stable for all $\theta$. However, the system also admits a limit cycle: for example, for $\theta = 200$ we find that for initial conditions (constant on the interval $[-\theta, 0]$) close to the steady state, the solution converges to $\bar{u}$. But for initial values far from the steady state, large-amplitude oscillations are seen (Fig. S8).
